# Systematic bias in malaria parasite relatedness estimation

**DOI:** 10.1101/2024.04.16.588675

**Authors:** Somya Mehra, Daniel E Neafsey, Michael White, Aimee R Taylor

## Abstract

Genetic studies of malaria parasites increasingly feature estimates of relatedness. However, various aspects of malaria parasite relatedness estimation are not fully understood. For example, estimates of relatedness based on whole-genome-sequence (WGS) data often exceed those based on more sparse data types. We explore systematic bias in relatedness estimation using theoretical, numerical and empirical approaches. Specifically, we use a non-ancestral model of pairwise relatedness to derive theoretical results; a simulation model of ancestry to independently verify and expand our theoretical results; and data on parasites sampled from Guyana to explore how theoretical and numerical results translate empirically. We show that allele frequencies encode, locus-by-locus, relatedness averaged over the set of sampled parasites used to compute them. These sample allele frequencies are typically plugged into the models used to estimate pairwise relatedness. Consequently, models of pairwise relatedness are misspecified and pairwise relatedness values are systematically underestimated. However, systematic underestimation can be viewed as population-relatedness calibration, i.e., a way of generating measures of relative relatedness. Systematic underestimation is unavoidable when relatedness is estimated assuming independence between genetic markers. It is mitigated when estimated using WGS data under a hidden Markov model (HMM), which exploits linkage between proximal markers. Estimates of absolute relatedness generated under a HMM using relatively sparse data should be treated with caution because the extent to which underestimation is mitigated is unknowable. That said, analyses dependent on absolute values and high relatedness thresholds are relatively robust. In summary, practitioners have two options: resolve to use relative relatedness estimated under independence or try to estimate absolute relatedness under a HMM. We propose various practical tools to help practitioners evaluate their situation on a case-by-case basis.

**Author summary:** Malaria genomic epidemiology is increasingly recognised as a tool for public health. Relatedness, which captures likeness derived from common ancestry, is a useful concept for malaria parasites. Analyses of malaria parasite relatedness are important for generating results on spatiotemporal scales relevant to disease control. Since shared ancestry is unobservable, relatedness must be estimated under a statistical model. However, not all aspects of malaria parasite estimation are fully understood, including the effects of different data types. In this work, we characterise systematic biases in estimates of malaria parasite relatedness. Our analysis is three-fold: we mathematically interrogate a non-ancestral model of relatedness to derive theoretical results; simulate parasite ancestries from first principles to yield numerical results; and perform an empirical case study of parasites sampled from Guyana. We show that bias may be particularly pronounced when using sparse marker data from inbred parasite populations, which are often found in pre-elimination settings. We chart out a practical roadmap to enable practitioners to assess epidemiological settings on a case-by-case basis. Our findings are relevant to applications in malaria genomic epidemiology that use relatedness directly or indirectly, including molecular surveillance and the genetic-based classification of treatment failure.

## 1 Introduction

Relatedness is a genome-wide measure of identity-by-descent (IBD); i.e., identity due to common ancestry [1, 2]. IBD is a useful concept when studying malaria-causing *Plasmodium* parasites because they sexually recombine [3]. Although obligate, recombination is only effective when genetically distinct parasites recombine. The probability that genetically distinct parasites recombine is context specific: it depends on parasite diversity, on the prevalence of infected hosts, and on the within-host diversity and prevalence of polyclonal infections among infected hosts [4]. As such, relatedness analyses reflect the recent, context-specific history of malaria parasites, generating epidemiologically relevant insight. For example, analyses of relatedness have been used to evaluate and inform efforts to reduce transmission [5, 6, 7], to elucidate population connectivity on granular spatiotemporal scales [8, 9, 10, 11, 12], to characterise the structure of inbred populations [13, 14], to resolve transmisson heterogeneity [15], and to identify regions of the parasite genome subject to recent selective pressure [14, 16, 17]. Relatedness has further applications in clinical trials of antimalarial drugs, i.e., in the classification of *Plasmodium falciparum* reinfection and recrudescence [18] and of *Plasmodium vivax* reinfection, recrudescence and relapse [19, 20].

IBD is not observable. As such, relatedness must be estimated under a statistical model. Various software exist for estimating relatedness between malaria parasites [16, 21, 22, 23, 24]. However, not all aspects of malaria parasite relatedness estimation are fully understood, especially those pertaining to systematic bias. For example, relatedness estimates based on dense whole-genome-sequence (WGS) data often exceed those based on sparse data, with striking zero-inflation of the sparse-data estimates (e.g., see Figure S2Q of [9]). As another example, consider the allele frequencies that are typically plugged into the models used to estimate relatedness between parasites. They are computed from a sample of parasites assuming inter-parasite independence — an assumption that contradicts pairwise relatedness estimation. They are thus liable to systematically bias relatedness estimates [9, 25, 26, 27, 28, 29]. Weir and Goudet [29] and others have addressed the consequent notion that relatedness is estimated relative to the average relatedness within the sample from which allele frequencies are computed. Implications of this construction have been addressed in the context of forensic typing [30], and case-control studies to characterise genetic determinants of human disease [31, 32, 33]. However, for practitioners of malaria genomic epidemiology, practical tools and guidelines that address these biases are, to the best of our knowledge, unavailable.

Here, we characterise systematic biases in malaria parasite relatedness estimation using three complementary approaches. First, we analyse theoretically non-ancestral models of pairwise relatedness, characterising the ramifications of the use of sample allele frequencies, shedding light on the aforementioned zero-inflation, and establishing common ground with Weir and Goudet [29]. Second, we simulate malaria parasite ancestries over successive generations of inbreeding, verifying independently our theoretical results, and elucidating systematic differences in relatedness estimates that are generated under models that assume inter-marker independence versus marker linkage due to proximity. Our numerical results help to explain differences between estimates based on sparse and dense data. Finally, using *P. falciparum* data from a highly-inbred parasite population in Guyana, we illustrate how our results based on theory and simulation translate empirically and the practical utility of various diagnostics. Beyond malaria genomic epidemiology, our results generalise to systems of largely haploid recombining eukaryotes [34, 35, 36, 37, 38] or highly-inbred diploid populations for which the haploid model of Leutenegger et al. [39] is applicable.

## 2 Methods

We characterise systematic biases in malaria parasite relatedness estimation using three complementary and independent approaches: by analysing statistical models of pairwise relatedness, by analysing simulated data with known parasite ancestries, and by analysing *P. falciparum* data from Guyana and Colombia. A summary of notation used throughout our main text is provided in Table 1. A detailed description of the methods described below is available in Appendices A and B, while a glossary of terms is provided in Appendix C.

**Table 1:** Summary of terms and notation.

| Terms / notation | Working definition |
| --- | --- |
| $\overline{\text{IBS}}_\ell$ | Locuswise average IBS sharing: the proportion of pairs in a sample (including self-self comparisons) that are IBS at a given marker $\ell$ |
| $\overline{\text{IBD}}_\ell$ | Locuswise average IBD sharing: the proportion of pairs in a sample (including self-self comparisons) that are IBD at a given marker $\ell$ |
| $\text{mean}(\overline{\text{IBD}}_\ell)$ | Locuswise average IBD sharing averaged over all markers $\ell = 1, \dots$ |
| $r$ | Relatedness parameter governing the marginal probability of IBD at each marker under non-ancestral models of pairwise relatedness |
| $\hat{r}$ | Maximum likelihood estimate of the parameter $r$; an estimate of pairwise relatedness |
| realised relatedness | Fraction of polymorphic markers across the genome that are IBD for a parasite pair |

### 2.1 Statistical models of pairwise relatedness

We derive theoretical results and estimate pairwise relatedness using models of relatedness that couple a non-ancestral model of latent IBD and non-IBD (nIBD) states describing a pair of parasites along a sequence of marker loci and a locuswise observation model.

The latent state model either assumes independence between markers or accounts for marker linkage under the intuition that IBD segments are fragmented at randomly-distributed recombination breakpoints over successive generations, with a genome-wide-constant recombination rate [39, 40, 41]. Of key interest is the pairwise relatedness parameter *r*, which describes the marginal probability that a given marker locus is IBD for a parasite pair, i.e., if we consider a single locus in isolation under the model then, 𝕡(IBD) = *r*.

The locuswise observation model provides an explicit link between latent and observable states. Observable states are either pairs of alleles or descriptives of identity-by-state (IBS); i.e., IBS for a pair of identical alleles and non-IBS (nIBS) otherwise. For IBS descriptives, the observational model takes the form 𝕡_obs_(IBS|IBD) = 1 in the absence of genotyping error (since IBD necessarily implies IBS) while 𝕡_obs_(IBS|nIBD) must be defined appropriately.

Under the coupled model of relatedness, the marginal likelihood of IBS at a given marker can be written

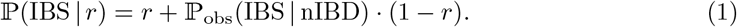

Here, we focus on maximum likelihood estimates 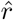 as an estimate of relatedness between a pair of malaria parasites. We do not address the estimation or identification of IBD segments.

It is standard practice to compute allele frequencies from data on a set of sampled parasites (either before or after removing replicates of apparent clones) and plug them into observation models. We refer to these frequencies as sample allele frequencies and to the observation models into which they are plugged as standard practice observation models. Other terms we use include (n)IBD to refer to IBD and nIBD collectively, (n)IBS to refer to IBS and nIBS collectively, hidden Markov model (HMM) of relatedness to refer to models of relatedness that allow for linkage between proximal markers, and independence model of relatedness to refer to models of relatedness that assume marker loci are independent. We use the word model to refer to models of relatedness, the models within the models of relatedness (i.e., (n)IBD models and (n)IBD-to-observation models) and the models within the (n)IBD models and (n)IBD-to-observation models (e.g., IBD-to-allele model, nIBD-to-IBS model). We use the term fraction to indicate pairwise averages over loci across the genome, and proportion to indicate locuswise averages over parasites within a population.

Here, we are primarily concerned with bias stemming from standard practice observation models. As a theoretical comparator, we adopt a corrected model of (n)IBS descriptives with conceptual parallels to Weir and Goudet [29]. The corrected model is not available practically, but illustrates theoretically the partial encoding of population relatedness in the sums of squares of sample allele frequencies. Further details of the observation models are provided in the results section. For clarity, genotyping error is not modelled and theoretical derivations are restricted to the independence model of relatedness.

### 2.2 Simulation model of parasite ancestries

While our theoretical results are derived under a non-ancestral pairwise framework, we perform numerical analyses under an ancestrally-informed simulation of a parasite population. Although our simulation framework is highly-simplified and does not fully recapitulate epidemiological reality, it enables independent verification of our theoretical results and allows us to examine the consequences of marker linkage.

A detailed description of the simulation model is provided in Appendix B. In brief, it captures successive generations of inbreeding, with the ancestry of each simulated parasite formulated as a mosaic of a bygone outbred founder population. Its main assumptions are as follows.

- Sufficient temporal separation between ‘generation zero’ and a bygone outbred founder population to break down marker linkage, whereby the ancestry of each individual in generation zero is sampled uniformly over the set of all possible founder mosaics; this construction distributes ancient low-level background relatedness broadly across the generation zero population.
- Discrete, non-overlapping generations of inbreeding from generation zero onwards, modelled using a transitive relationship graph with sibling/clonal/stranger edges as per [20]; relationship graphs are obtained by amalgamating uniformly-sampled subgraphs of a fixed size [42], biasing the distribution towards small, balanced clusters of siblings and clones.
- Stochastic drift arising from the random crossing of parental genotypes from generation (*k −*1) to yield filial genotypes in generation *k*, with an augmented probability of selfing and sibling-sibling crosses serving as a proxy for monoclonal mosquito infection and serial cotransmission [43].
- The absence of immigration, mutation, selection and population structure under a small, fixed parasite population size.
- A genome-wide constant recombination rate, with markers treated as nominal point polymorphisms.

When we analyse simulated data, the ground truth against which we characterise systematic bias in pairwise relatedness estimates is the fraction of polymorphic marker loci IBD for a given parasite pair; otherwise known as realised relatedness [2]. There is a conceptual distinction between realised relatedness and the relatedness parameter *r* of the non-ancestral pairwise model: given a finite number of markers, a fixed value of *r* gives rise to a *distribution* of realised relatedness [2, 41]. However, since we model recombination from first principles under our simulation model, our simulation framework does not yield a direct analogue for the non-ancestral relatedness parameter *r*.

### 2.3 Case study of *P. falciparum* data from Guyana and Colombia

To show our theoretical and numerical results are practically relevant, we present a case study from a highly inbred parasite population. Specifically, we consider a set of high-quality isolates from clinical infections that are deemed monoclonal, collected in passively sampled symptomatic patients in Guyana between 2016–2020 [44] and Colombia between 1993–2017 [14], for which WGS data are available. Variants were called in accordance with best practices stipulated by GATK and the MalariaGEN Pf3k consortium [14, 44]. We restrict our attention to biallelic SNPs that are in core nuclear regions of the genome [45], and are polymorphic among the sampled infections, masking any variant calls that are heteroallelic or have read support below 5 (based on the DP tag). We remove variant sites with missingness *>* 30% across sampled infections and sampled infections with missingness *>* 30% across SNPs to yield a WGS SNP dataset comprising *n* = 306 sampled infections (*n* = 278 from Guyana, *n* = 28 from Colombia) and *n* = 30694 polymorphic sites. Results in the main text are restricted to the sample from Guyana, comprising *n* = 278 sampled infections and *n* = 16115 polymorphic biallelic SNPs. Sparse datasets are generated by down-sampling variant sites uniformly at random without replacement.

### 2.4 Data analyses

All data analyses are done in R [46]. Relatedness estimates based on allelic states are generated using the R package paneljudge [24], which implements the HMM of relatedness with allelic observations and its independent counterpart, and employs a maximum likelihood estimation scheme. The model of independent (n)IBD states with (n)IBS observations has been custom coded in R. We do not estimate relatedness using a HMM with (n)IBS observations. All code and a minimum analysis dataset for the empirical case study is available on GitHub: https://github.com/somyamehra/PlasmodiumRelatednessBias

## 3 Results

A detailed description of the results below can be found in Appendix A, which includes an overview of the support (theoretical, numerical, and/or empirical) per result (Table A.1).

### 3.1 Theoretical results

#### 3.1.1 Standard (n)IBD-to-observation models are twice misspecified

In standard practice, (n)IBD-to-observation models are informed by sample allele frequencies [9, 10, 16, 22, 41]. Taylor et al. [41] allude to potential misspecification arising from this construction: while relatedness estimation seeks to estimate dependence between individuals, the sample allele frequencies, and thus any observation models into which they are plugged, are constructed under the implicit assumption of independence between individuals. Weir and Goudet [29] have articulated the notion that relatedness is consequently estimated relative to the average relatedness within the sample.

Here, we argue that standard (n)IBD-to-observation models are twice misspecified:

1. The standard IBD-to-allele model, whereby the probability of an IBD pair exhibiting allele *q* is given by the sample frequency of allele *q*, is implicitly predicated on the independence of allelic and IBD states which may not hold in reality (Equation (A.10)), particularly in the presence of selection.
2. The standard nIBD-to-IBS model, under which the probability of an IBS pair given a nIBD pair is equal to the proportion of IBS pairs in the set of sampled parasites (calculated by taking the sums of squares of sample allele frequencies), is inflated by the encoding of average locuswise relatedness (Equation (A.12)). Weir and Goudet [29] similarly exploit IBS descriptives rather than allelic states for conceptual clarity.

Removing clonal replicates will not circumvent IBD-to-allele misspecification although it seems reasonable to expect it might mitigate it; otherwise, consequences of the IBD-to-allele misspecification are case-specific and beyond the scope of our current study. For the remainder of this study we focus on nIBD-to-IBS misspecification because it is more pervasive and has systematic consequences.

An illustration of nIBD-to-IBS misspecification is shown in Box 1, echoing the work of Weir and Goudet [29]. We can summarise this form of misspecification mathematically as follows. Denote by 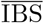 and 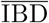 the proportion of pairs of sampled parasites (including self-self comparisons) that are IBS and IBD respectively at a given locus (Table 1, but with locus identifiers dropped for notational convenience). Under the standard nIBD-to-IBS model, we set

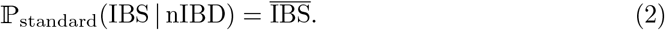

A proportion of IBS sharing, however, is attributable to IBD. To adjust for locuswise relateness in the set of sampled parasites, we would need to average IBS sharing over nIBD pairs only, that is,

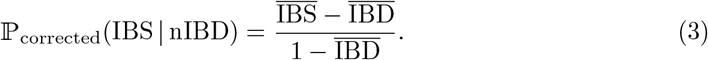

The correction (Equation (3)) cannot be implemented in practice because (n)IBD states are unobservable. However, it provides a theoretical basis for understanding the misspecification of the standard nIBD-to-IBS model. To see how relatedness structure, in the form of locuswise average relatedness 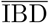, is implicitly embedded in the standard nIBD-to-IBS model, we rearrange Equation (3) to yield

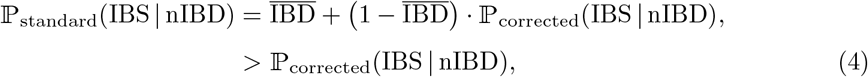

where the strict inequality is just a technicality. Since we have included self-self comparisons (i.e., comparisons with replacement) for consistency with standard nIBD-to-allele models [16, 21], 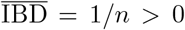 for an outbred sample of size *n*. The finite sample adjustment of [47], which considers pairwise comparisons without replacement, yields 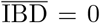 for an outbred sample. However, under either construction, the form of the theoretical correction to the nIBD-to-IBS model is identical.

In summary, the probability of IBS sharing for nIBD pairs is systematically overestimated under the standard nIBD-to-IBS model, with a particularly pronounced effect in inbred populations with large 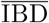.

##### Box 1

**Illustrating misspecification of the standard nIBD-to-IBS model**

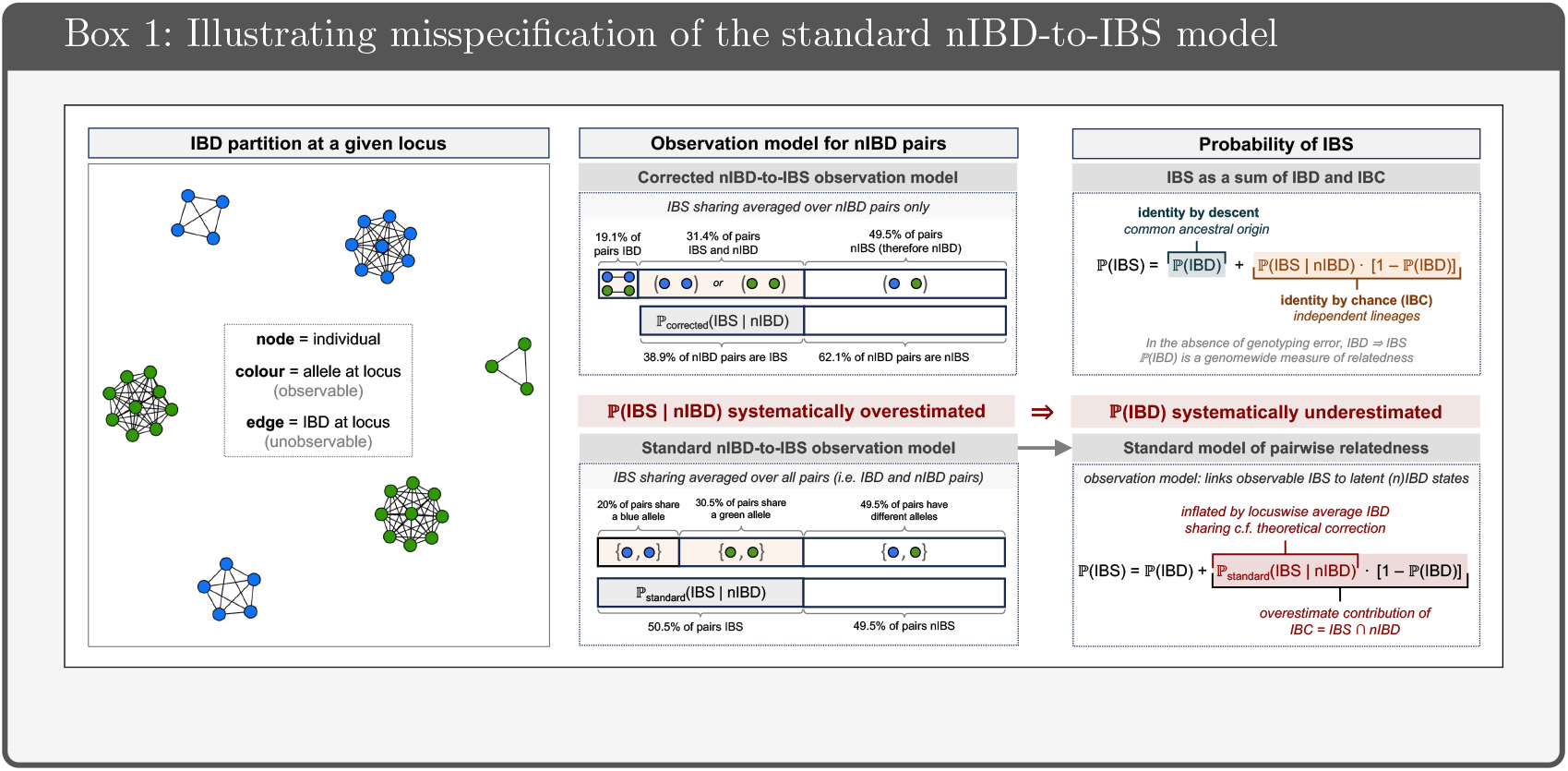

###### Standard nIBD-to-IBS model

At a given locus, *b* = 17 blue nodes and *g* = 21 green nodes are observed. The proportion of pairs that are IBS, which we denote 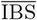, can be written as the sum of squares of the blue *b/*(*b* + *g*) and green *g/*(*b* + *g*) allele frequencies:

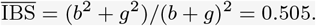

Under the standard nIBD-to-IBS model the probability of an IBS pair given an nIBD pair is equal to the overall proportion of pairs that are IBS:

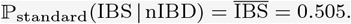

###### Corrected nIBD-to-IBS model

There are three green clusters in the IBD partition, of size *c*_1_ = 9, *c*_2_ = 9 and *c*_3_ = 3. Out of a total of (*b* + *g*)^2^ = 1444 possible pairwise comparisons, this means that

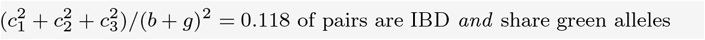

Likewise, since there are three blue clusters of size *s*_1_ = 8, *s*_2_ = 5, *s*_3_ = 4

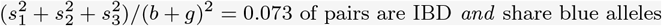

The proportion of pairs which are IBD is then

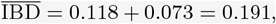

while the proportion of pairs that are IBS and nIBD is

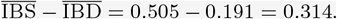

To construct a model of IBS specifically for nIBD pairs, we would ideally focus on IBS sharing in the proportion of 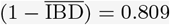 pairs that are nIBD, that is,

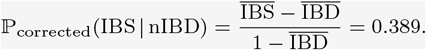

###### Misspecification of the standard n IBD-to-IBS model

In failing to adjust for IBD sharing under the standard nIBD-to-IBS model, we have *overestimated* the probability of IBS sharing for nIBD pairs:

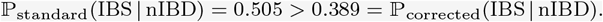

#### 3.1.2 Misspecification: relative and zero-valued relatedness estimates

Here, we examine systematic bias in maximum likelihood estimates 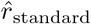 generated under the standard nIBD-to-IBS model (Equation (2)), relative to hypothetical estimates 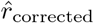 generated under the corrected nIBD-to-IBS model (Equation (3)). For conceptual clarity, we enforce the assumption of marker independence (consequences of marker linkage are explored in our numerical analyses, Section 3.2).

We can intuit that misspecification of the standard n IBD-to-IBS model will lead to systematic underestimation of pairwise relatedness, that is, 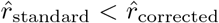(Box 1). Two additional consequences for which we have theoretical support are as follows.

1. The standard pairwise relatedness parameter *r*_standard_ can be re-interpted as a relative measure of deviation from population-averaged locuswise relatedness (Appendix A.2.2.1). The marginal likelihood of IBS sharing at a given marker *ℓ* under the standard model with relatedness parameter *r*_standard_ is equivalent to the likelihood under the corrected model with relatedness parameter 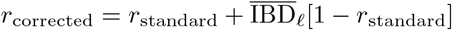, where 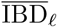 is the proportion of sampled parasites that are IBD at locus *ℓ*, that is, 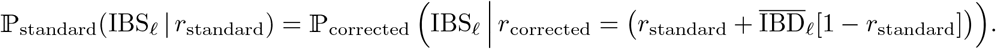 In practice, the average locuswise relatedness 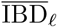 will vary across loci *ℓ*. In a hypothetical population with identical average locuswise relatedness over all loci, 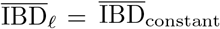, we obtain the functional relationship 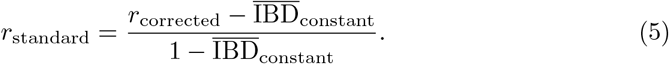 This interpretation of relative relatedness echoes the work of Weir and Goudet [29].
2. Relatedness estimates can be stratified by average sample relatedness: zero below, positive otherwise (Appendix A.2.2.2). To understand why this is the case, using Equations (1) and (4), we observe that 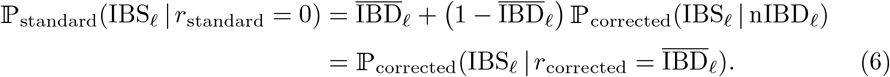 That is, plugging *r*_standard_ = 0 into the standard model likelihood of IBS at a given marker *ℓ* is equivalent to plugging in the average locuswise relatedness 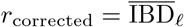 into the corrected model likelihood. In other words, population-averaged locuswise relatedness is implicitly encoded in the standard model even in the case *r*_standard_ = 0, and parasite pairs with less IBS sharing than predicted under population-averaged relatedness (given explicitly by the threshold (A.25)) are assigned zero estimates 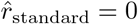. Similar observations have been made previously by Weir and Goudet [29].

Accounting for variability in 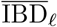 across loci *ℓ*, we predict estimates 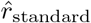 against the theoretical comparator 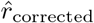 to exhibit a fuzzy elbow-like characteristic, with a change point in the vicinity of the mean locuswise average IBD sharing, mean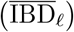 (Figure 1A). If the distribution of 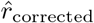 is positively skewed, we expect pronounced zero inflation in estimates of *r*_standard_ under the standard nIBD-to-IBS model.

**Figure 1:**
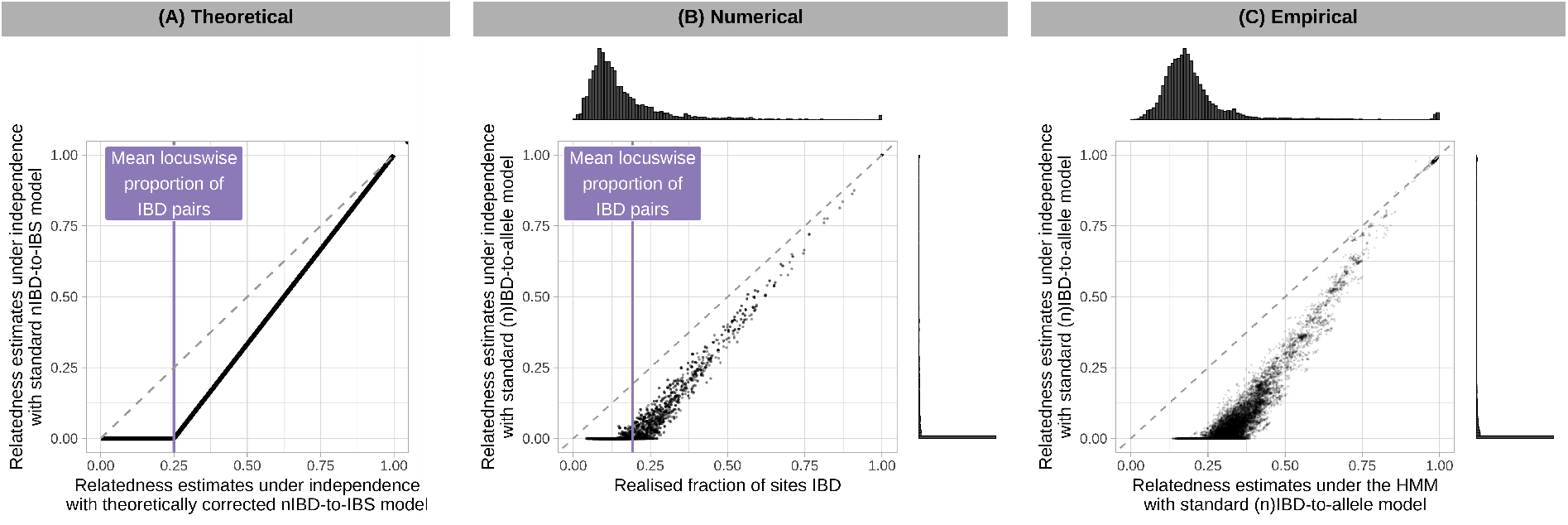
Characterisation of systematic bias in pairwise relatedness under (n)IBD independence. We characterise bias against three comparators: (A) 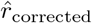 estimates computed theoretically; (B) realised relatedness computed using simulated data; and (C) 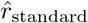 estimates computed using the HMM and WGS data on *P. falciparum* from Guyana. In each case, we recover an elbow-like characteristic with change point near the mean locuswise proportion of IBD pairs mean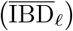.

### 3.2 Numerical results

#### 3.2.1 Theoretical validation

Relatedness estimates generated under the independence model of relatedness using simulated data support theoretical results as follows. In line with standard nIBD-to-IBS model misspecification, systematic bias in estimates of *r*_standard_ increases as mean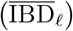 increases (Appendix A.3.1). In line with the proposed nIBD-to-IBS model correction, 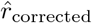 are largely unbiased (Appendix A.3.2, Figure 2A vs 2C). The realisation of bias-mitigation under (n)IBD independence supports the notion that systematic bias of 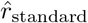 can be attributed to the partial encoding of sample relatedness in standard observation models. In line with expected zero-valued estimates, 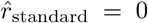 for simulated pairs that exhibit a smaller fraction of IBS markers than that which is expected for *r*_standard_ = 0 under the standard nIBD-to-observation model (Appendix A.3.4). In line with Figure 1A, plots of 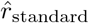 against realised relatedness yield an elbow-like characteristic, branching approximately at mean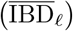 (Figure 1B).

**Figure 2:**
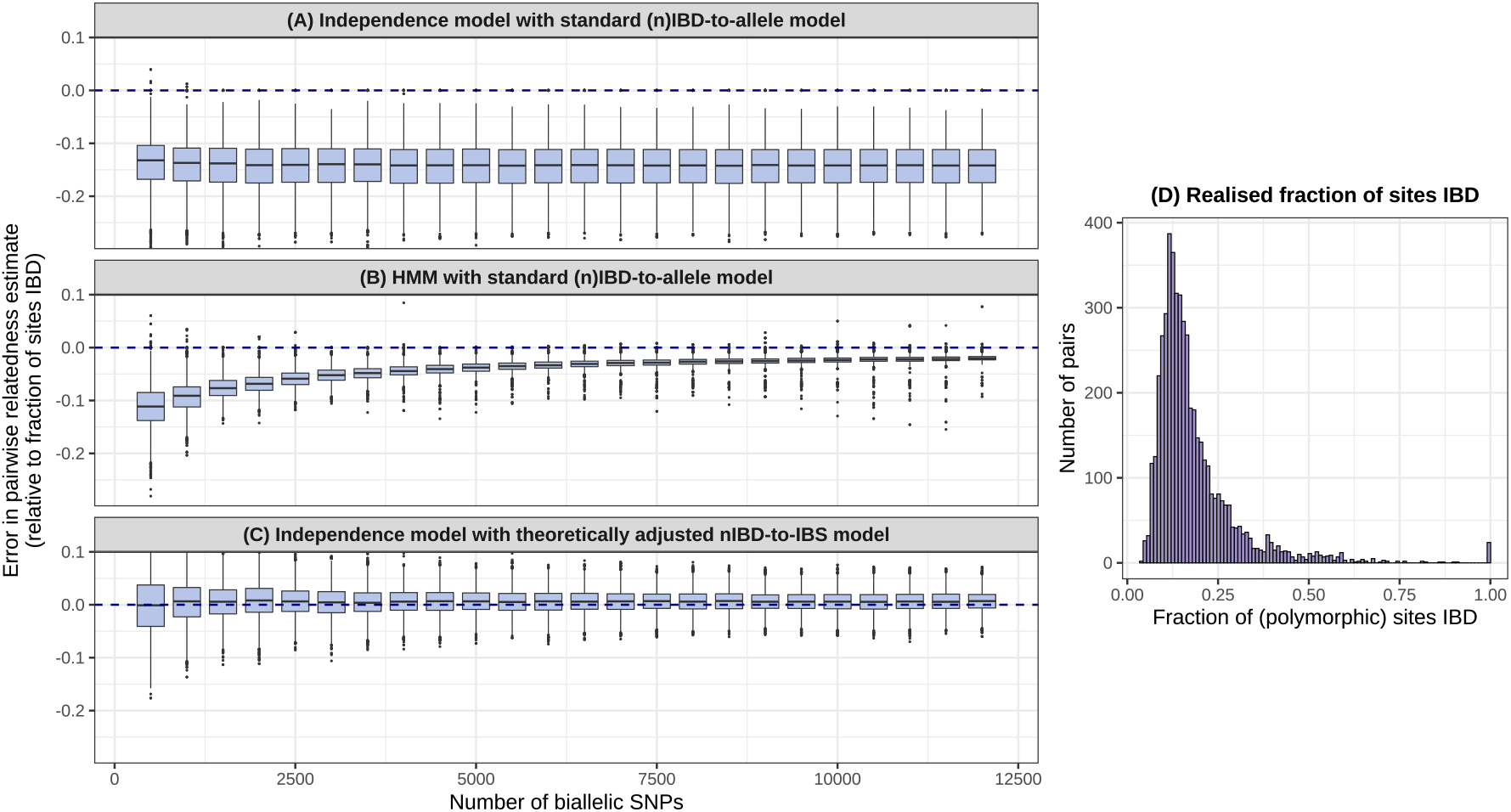
Summary of pairwise relatedness for simulated data after 10 generations of inbreeding. Systematic bias is shown as a function of marker density, with the fraction of (polymorphic) marker loci IBD taken as the ground truth, for (A): 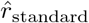 under (n)IBD independence and the standard nIBD-to-allele model; (B) 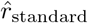 computed using the HMM and the standard nIBD-to-allele model; (C): 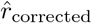 under (n)IBD independence and the corrected nIBD-to-IBS model. (D): Histogram of realised relatedness (fraction of polymorphic sites that are IBD for simulated parasites).

#### 3.2.2 Exploiting marker linkage can mitigate bias

While systematic underestimation given (n)IBD independence persists irrespective of marker density (Figure 2A), it is mitigated as a function of marker density when data are analysed under the HMM (Figure 2B). We attribute this trend to the exploitation of increasingly detailed linkage information under the HMM, reducing the reliance of *r*_standard_ estimation on the standard observation model, which is misspecified. We thus propose a dense data diagnostic that leverages increased precision under the HMM (Appendix A.3.5): for dense data, comparison of 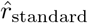 estimates under (n)IBD independence vs the HMM is expected to yield an elbow-like characteristic (analogous to Figure 1B), which can be used to ascertain both the severity of underestimation and the approximate degree of relatedness mean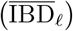 averaged over the set of sampled parasites. Zero inflation renders mean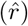 under (n)IBD independence a poor approximation of mean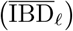.

For sparse data that do not encode linkage information, analyses of simulated data under the independence model of relatedness confirm that bias-mitigation would be possible if model-adjustment were available, and that model-adjustment would be sufficient (Figure 2C).

#### 3.2.3 Relative vs absolute relatedness

Across successive generations of inbreeding, realised relatedness between siblings (gray shading, Figure 3) is systematically enriched above 0.5. For dense data, we can interpret 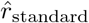 under the HMM (navy blue, Figure 3) as a measure of absolute relatedness which recapitulates this enrichment. In contrast, 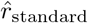 for siblings under (n)IBD independence (orange, Figure 3) is consistently centred around 0.5, supporting reinterpretation as a measure of relative relatedness. This points towards two choices for practitioners: the analysis of absolute relatedness under a HMM, which requires sufficiently dense genotypic data; or the use of relative relatedness under (n)IBD independence, which warrants careful interpretation in cross-population comparisons (the degree of bias may vary across transmission settings, but relative relatedness may have utility, for instance, as an approximate relationship proxy).

**Figure 3:**
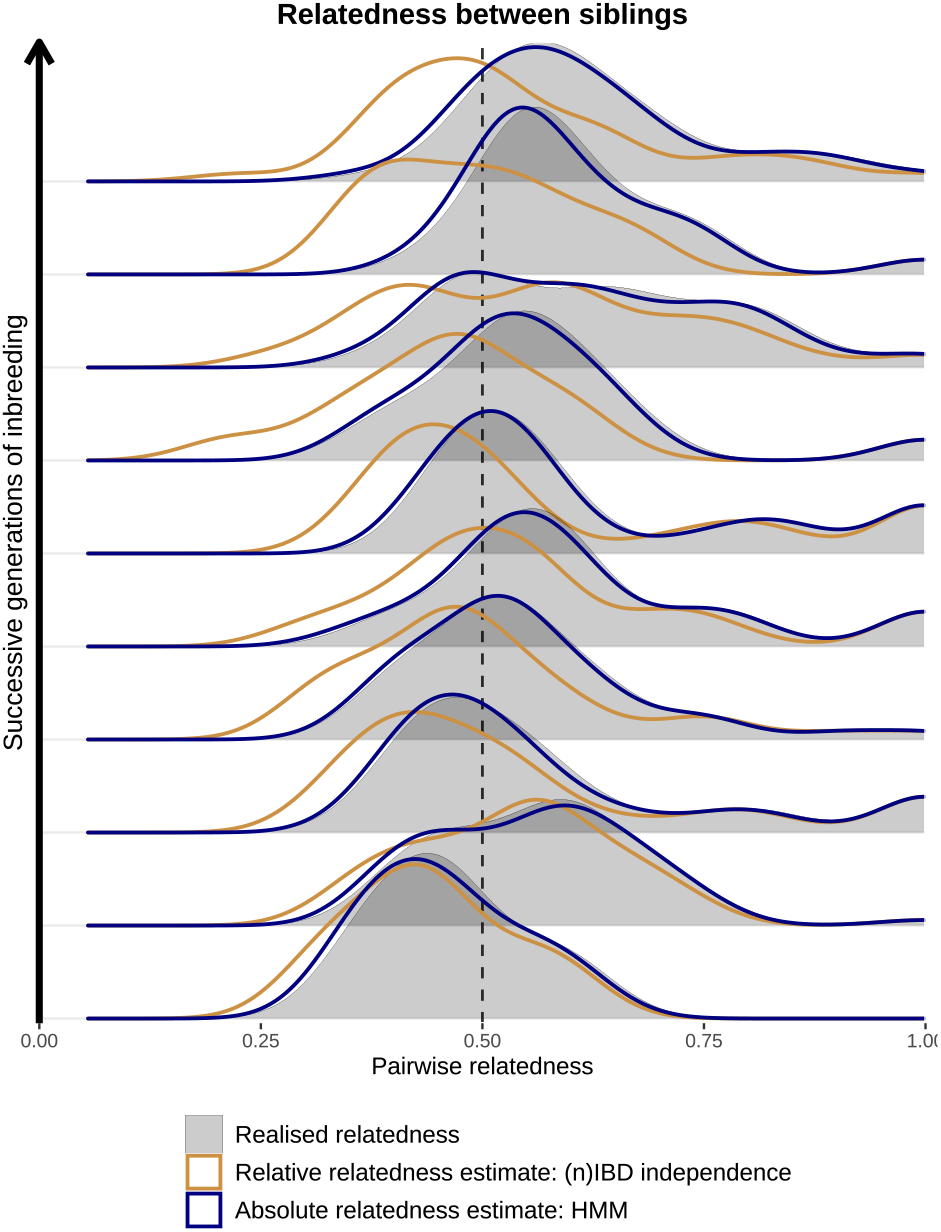
Summary of pairwise relatedness for simulated siblings across successive generations of inbreeding. We compare realised relatedness (i.e., the fraction of (polymorphic) sites that are IBD for a given parasite pair) against 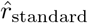 predicated on the standard nIBD-to-allele model with (n)IBD independence (relative relatedness estimates) versus the HMM (absolute relatedness estimates).

### 3.3 Empirical results

#### 3.3.1 Case study: inbred parasite population from Guyana

To illustrate the practical implications of our findings, we analyse WGS data from *n* = 278 high-quality *P. falciparum* isolates (deemed to be monoclonal) sampled from Guyana in 2016–2020.

Based on our numerical results (specifically, Figure 2B), estimates of relatedness generated under the HMM using WGS data are expected to exhibit relatively little bias and therefore serve as a pragmatic gold-standard. To gauge the severity of systematic underestimation due to standard observation model misspecification, we draw on the dense data diagnostic proposed in Section 3.2.2: a comparative plot of dense-data 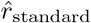 estimates generated under (n)IBD independence model and HMM (Figure 1C). The position of the change point suggests that the mean locuswise proportion of IBD pairs is in the vicinity of mean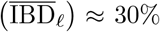. Across parasite pairs, the average estimate 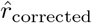 under the HMM using WGS data is 30.4%.

In addition to systematic bias stemming from model misspecification, uncertainty due to marker sparsity may become significant for sparse data [10]. We expect the point at which uncertainty/variance masks the elbow-like characteristic to be dependent on the average relatedness within the set of sampled parasites. A sparse-dense data diagnostic, comprising comparative plots of estimates generated under (n)IBD independence using sparse data vs estimates generated under the HMM using dense data, can be used to elucidate this trade-off because under (n)IBD independence, increasing marker density does not mitigate systematic bias but does reduce uncertainty in pairwise relatedness estimates. For 278 *P. falciparum* isolates from Guyana, Figure 4A suggests bias due to elevated population relatedness dominates uncertainty due to marker sparsity: even at low marker densities, systematic underestimation is apparent.

**Figure 4:**
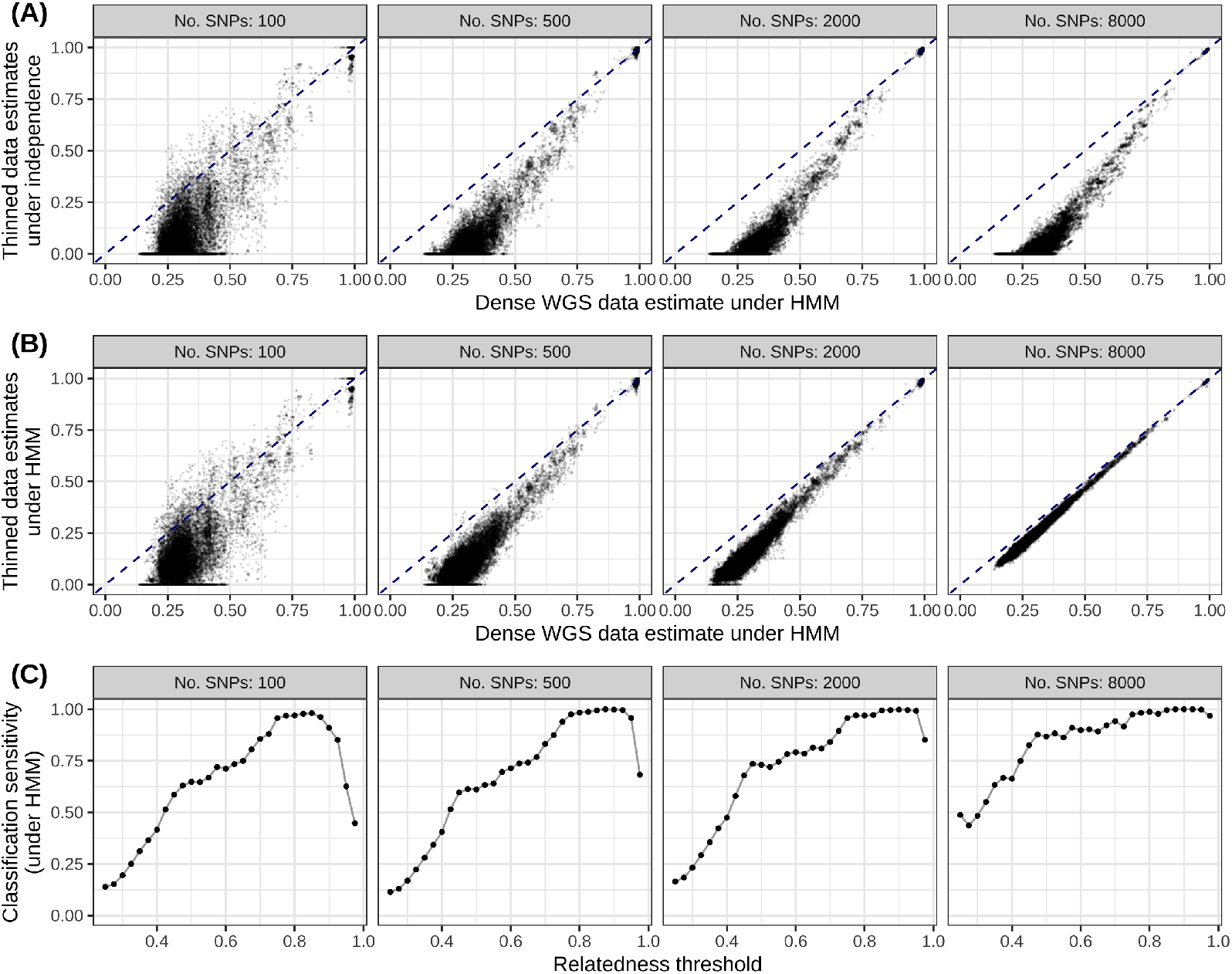
Consequences of marker sparsity on pairwise relatedness estimates for inbred parasites sampled from Guyana. Since IBD is unobservable, we treat relatedness estimates generated under the HMM for the complete dense WGS SNP dataset as a truth value. Thinned data have been generated by downsampling uniformly at random without replacement biallelic SNPs that are polymorphic among the 278 samples. ===== (A) Relatedness estimates generated under (n)IBD independence with thinned data vs the HMM with dense data. ===== (B) Relatedness estimates generated under the HMM with thinned data vs dense data. ===== (C) Sensitivity of thinned data for classifying related parasite pairs, when analysed under the HMM of relatedness. For each IBD threshold *r*_threshold_, we record a true positive (TP) if estimates generated under the HMM for both thinned and dense data lie above *r*_threshold_; and a false negative (FN) if only the HMM estimate for dense data lies above *r*_threshold_. The sensitivity is given by the ratio TP/(TP+FN). ===== All estimates are generated using the standard (n)IBD-to-allele model.

A possible strategy for offsetting systematic bias is to exploit linkage information under the HMM regardless of marker density. Analysing sparse data under the HMM, however, yields an intermediary regime, whereby systematic underestimation driven by nIBD-to-IBS misspecification is only partially mitigated (Figure 4B). Unlike estimates under the indepen-dence model — which can be re-interpreted as relative measures of relatedness, intrinsically calibrated for locuswise average relatedness within a set of sampled parasites (Section 3.1.2) — sparse data HMM estimates do not have a clear interpretation.

The error structure in sparse data HMM estimates is heteroskedastic: underestimation is most pronounced for parasite pairs with low relatedness. At the other end of the spectrum, genomic epidemiology applications typically focus on highly related parasite pairs, often using a threshold-based approach [9, 14, 16, 48, 49, 50]. Since relatedness is systematically underestimated using sparse data, false positives (pairs with low levels of relatedness that appear highly related) are relatively rare. The sensitivity of sparse data (analysed under the HMM) in identifying parasite pairs above a relatedness threshold *r*_threshold_ is shown in Figure 4C. When relatedness thresholds are sufficiently high, sparse marker data in the intermediary regime suffice; sensitive classification for low relatedness thresholds, however, requires higher resolution data [50]. We are reluctant to posit marker density thresholds necessary to sensitively identify highly-related parasite pairs in generality, because the degree of linkage structure within a set of sampled parasites depends on the distribution of shared IBD segment lengths — which, in turn, is driven by demographic processes that we neither fully understand, nor control. The downsampling of WGS data on a case-by-case basis, however, may provide setting-specific insights.

#### 3.3.2 Population structure

Since our theoretical and numerical results assume a single parasite population, we additionally perform a joint analysis of the *n* = 278 isolates from Guyana and *n* = 28 isolates from Colombia. Principal coordinates analysis (PCoA) of pairwise fractions of IBS markers yields two distinct clusters, stratified by country (Figure A.12). Two diagnostics — namely, a multimodal distribution of pairwise fractions of IBS markers, and systematic differences between relatedness estimates under the standard nIBD-to-IBS vs (n)IBD-to-allele model — corroborate the presence of population structure (Appendix A.4.1).

In this setting, the dense data diagnostic is characterised by multiple elbows, corresponding to different within- and across-population comparisons (Figure A.15). Given sample allele frequencies constitute a weighted average across subpopulations, the interpretation of the corresponding branch points is unclear; an observation model predicated on subpopulation-stratified allele frequencies [21], may yield more interpretable results. Further discussion is provided in Appendix A.4.

## 4. Discussion

### 4.1 Summary and interpretation of results

We have characterised systematic biases in pairwise relatedness estimates, providing theoretical, numerical and/or empirical support for our results (Table A.1) via a three-fold approach (interrogation of non-ancestral models of pairwise relatedness; simulation under an ancestrally-informed model of population relatedness; and a case study of *P. falciparum* WGS data).

Our results are motivated by but not limited to analyses of malaria parasites. They extend to any system concerned with pairwise relatedness of predominately haploid recombining eukaryotes (e.g., *Cryptosporidium hominis* [37] and *Cryptosporidium parvum* [36], leading causes of human and zoonotic cryptosporidiosis respectively; *Coccidioides* species which give rise to human coccidioidomycosis [34]; *Cryphonectria parasitica*, the pathogenic agent responsible for Chestnut blight [35] and *Marchantia polymorpha*, a model species of liverwort [38]) or highly-inbred populations of diploid organisms for which pairwise relatedness can be interrogated using a haploid model [39].

Misspecification of standard (n)IBD-to-observation models, which are predicated on sample allele frequencies, constitutes our conceptual starting point [25, 29, 41]. Theoretically, we demonstrate via the nIBD-to-IBS model that the implicit embedding of population-averaged locuswise relatedness in standard observation models is pervasive and can lead to the systematic underestimation of pairwise relatedness. Our theoretical results bear strong conceptual similarity to the work of Weir and Goudet [29]. Both theoretical and numerical analyses support the re-interpretation of pairwise relatedness under the independence model as a relative measure: non-zero estimates are calibrated intrinsically for relatedness averaged over the set of sampled parasites, while zero estimates are non-informative and flag parasite pairs with below-average relatedness, echoing the work of Weir and Goudet [29]. By introducing linkage structure within the simulation model, we show numerically that the exploitation of linkage structure using a HMM can mitigate bias when genotypic data are sufficiently dense. We thus propose a dense data diagnostic (i.e., a plot of pairwise relatedness estimates generated using dense data under independence versus a HMM) to elucidate average sample relatedness and the potential severity of systematic bias. We illustrate the use of the dense-data dignostic using *P. falciparum* data. Also using *P. falciparum* data, we characterise the consequences of marker sparsity by downsampling markers. We find that the analysis of sparse data under an HMM may yield an intermediary regime that is difficult to interpret because systematic underestimation is only partially mitigated, but that threshold-based analyses are relatively robust.

The functional consequences of our findings are heavily context-dependent: the severity of systematic bias depends on average sample relatedness, which is contingent on the history of effective recombination in the parasite population; while mitigation of bias under the HMM is a function of linkage structure, which is largely governed by recent ancestry and the length of shared IBD segments. Rather than issuing a set of concrete rules, we chart out a roadmap — starting with representative WGS data from a parasite population of interest — for practitioners to evaluate specific settings on a case-by-case basis. Firstly, to assess whether the setting of interest is one where elevated population relatedness warrants caution around the estimation and interpretation of pairwise relatedness, we propose the use of the dense data diagnostic. If systematic bias is found to be pronounced, we suggest a heuristic: downsampling loci. A sparse-dense data diagnostic, comparing estimates for downsampled data under the independence model against gold-standard WGS estimates under a HMM, can elucidate the relative importance of systematic bias at low marker densities where estimates are inherently more uncertain due to limited marker informativity. Comparison of downsampled vs WGS estimates under a HMM can guide the use of a HMM (which partially mitigates bias) or the independence model (which yields interpretable bias) at a particular marker density in the setting of interest. Examination of the heteroskedastic error structure can also facilitate the identification of thresholds above which highly-related parasite pairs can be identified sensitively. Analytic choices will plausibly vary across transmission settings and use cases.

Using data simulated under the (non-ancestral) standard HMM model of relatedness, relatively accurate relatedness estimates were generated using moderate marker counts [41]. That was a well specified setting: data were simulated under the standard HMM model. In this study we consider misspecification of the standard HMM model. In a highly-inbred setting, bias-mitigation due to misspecification of the standard HMM model requires dense data. For sparse data in isolation, relatedness estimates can either be characterised using high relatedness thresholds or generated under (n)IBD independence and interpreted relatively.

Selection can likewise modulate IBD segment length distributions which, in turn, can have a bearing on estimates of genome-wide IBD [51]. For example, Guo et al. [51] have shown that positive selection can also bias relatedness estimation. However, they find selection-bias is minor when background relatedness is high. Taken together, our study and that of Guo et al. [51] suggest selection bias dominants when transmission is high; bias due to observation model misspecification dominates when transmission is low.

### 4.2 Limitations

#### IBS descriptives

The theoretical re-interpretation of pairwise relatedness under independence as a relative measure is predicated on IBS descriptives, as in Weir and Goudet [29]. In practice, pairwise relatedness is estimated using allelic states rather than IBS descriptives [10, 16, 21, 22, 24, 41]. Misspecification of the IBD-to-allele model — arising from the non-independence of IBD and allelic states — is sensitive to the unobservable relatedness structure at each locus. It is not necessarily rectified by removing replicates of clonal parasites, and may be particularly pronounced in the presence of selection [51]. As such, there may be systematic differences between pairwise relatedness estimates based on IBS descriptives versus allelic states. A rigorous examination of the latter would potentially require joint estimation of relatedness across a set of sampled parasites, which is beyond the scope of the current manuscript.

#### Reliance on dense genotypic data

The proposed diagnostics for gauging average sample relatedness and the severity of systematic bias in pairwise relatedness estimates rely on dense data from a representative population of interest. To mitigate systematic bias, we suggest leveraging linkage structure which necessarily mandates dense data for inbred parasite populations. In practical terms, dense data translates to WGS data. These data may not be readily available, or may be prohibitively expensive to generate [52, 53]. Sparse marker data, which are often of greater practical utility [52, 53], cannot be evaluated or corrected for systematic bias in isolation under our framework. For sparse data in isolation, we offer the re-interpretation of pairwise relatedness estimated under (n)IBD independence and, for relatedness estimated under the HMM, advocate caution to be exercised at low relatedness thresholds.

#### Demographic processes

Clearly, demographic processes have a large effect on real data but are neither explored theoretically nor numerically, to avoid overburdening the manuscript. The effects of selection and immigration on the underlying relatedness structure of a set of sampled parasites are not considered. Our ancestral simulation model is very simple and does not recapitulate epidemiological reality. Transmission dynamics — which govern the propensity for selfing versus inbreeding versus outbreeding — are not explicitly modelled. We do not account for fluctuations in parasite population sizes, for instance, due to bottlenecks or control interventions, which may be particularly relevant to the context in which genomic epidemiology studies are performed.

#### Population structure

While we propose several diagnostics to screen for population structure, we did not explore population structure theoretically or numerically [54]. Under the non-ancestral theoretical framework, locuswise average IBD sharing, IBD_*ℓ*_, is a representative summary statistic for a set of parasites sampled from the same population. In the presence of population structure, with variable levels of relatedness within and between populations, the predicted elbow-like characteristic of Figure 1 may break down; we posit the emergence of multiple elbows, corresponding to different cross-population comparisons (see Appendix A.4 for an empirical example). Zero-inflation may not be present for cross-population comparisons with lower relatedness than the cross-population average. The simulation model could be used to simulate different populations that unite and mix in order to interrogate the consequences of population structure. The simulation model of Guo et al. [51], which accounts for population structure, could be used as a complementary approach to our simulation model. The framework of Weir and Goudet [29], which jointly characterises relatedness and population structure, could also be drawn on. Theoretically, the cross-population model of pairwise relatedness provided by hmmIBD could be analysed [21].

#### Multi-allelic markers

We have derived results for multi-allelic markers in the absence of genotyping error, under the assumption that there are no systematic biological differences between multi-allelic and biallelic markers. That is to say, both are treated a nominal point polymorphisms, whose alleles can be modelled as categorical random variables [41]. We do not report our results on multi-allelic markers, because they are not meaningfully different to biallelic markers under these simplistic assumptions. In reality, the ancestral processes governing biallelic markers and multi-allelic markers likely differ. Multi-allelic markers additionally possess an ordinal genotyping error structure that is overlooked in current methods commonly used to estimate relatedness between malaria parasites [16, 21]. The practical ramifications could be explored using empirical data: qualitatively compare graphs of relatedness (as in [16]), where relatedness is estimated using multi-allelic markers (e.g., microhaplotypes) versus biallelic markers (e.g., SNPs) of equal informativeness (i.e., markers sets whose composite score of average effective cardinality multiplied by marker count is the same; see [41]).

#### Data sparsity under simulation

We explore data sparsity using real data only, not simulated data. The simulation model is not constructed in such a way that lends itself to the exploration of the effect of data sparsity. More specifically, under the simulation model, we intentionally elevate the recombination rate to compensate for computational constraints (small population size, few generations, single chromosome) and simplifying assumptions (no immigration, no mutation). By way of comparison, we simulate 24000 equidistant markers along a single chromosome of length 48 Morgans, while the *P. falciparum* nuclear genome comprises 14 choromosome with cumulative length 17 Morgans [45]. 24000 markers is larger than the *∼* 15000 SNPs reported by Miles et al. [45] in genetic crosses of *P. falciparum*, but smaller than the number of SNPs and indels combined (*∼* 28000); the simulated inter-marker distance of 0.002 Morgans is an order of magnitude larger than 0.0002 Morgans based on these crosses. Elevating the recombination rate generates simulated data whose estimates of relatedness resemble data; however, this also leads to more IBD segments among simulated data which, due to the law of large numbers, offsets the effect of data sparsity. An exceedingly large recombination rate, however, yields little linkage structure, whereby bias is only partially mitigated under the standard HMM model, even for dense data.

### 4.3 Future work

#### Generating community resources

The dense-data elbow-like plot, which can be used to diagnose the extent of underestimation, requires WGS data, which are often financially prohibitive [52, 53]. As a practical resource for the malaria genomic epidemiological community, catalogues of dense-data elbow-like plots could be generated for published WGS datasets [55]. Doing so would first require partitioning of samples into sets for which population structure is not a complicating factor. This exceeds the scope of this study. It is feasible, however, using, for example, PCoA and the diagnostics we propose in Appendix A.4.1.

#### Recurrent infection classification in therapeutic efficacy studies

Systematically elevated population relatedness is liable to impair the genetic resolution of recurrent infections during therapeutic efficacy studies in low transmission settings. How best to deal with elevated population relatedness in recurrent classification is not yet understood. One approach involves embedding a relatedness inflation factor into a classification model [20]. However, our results suggest the inclusion of inflation factors in classification models that use sample allele frequencies may be unnecessary because sample allele frequencies already partially encode population-averaged locuswise relatedness.

#### Analyses using confidence intervals

A key consideration for sparse genotypic data is the significance of systematic bias stemming from model misspecification in the presence of uncertainty attributable to marker sparsity. While we suggest downsampling loci to evaluate this trade-off heuristically, the incorporation of confidence intervals, as recommended in [41], may yield a more statistically-principled way to evaluate this trade-off.

#### Algorithmic correction for population-averaged locuswise relatedness

Unbiased relatedness estimation using sparse marker data requires joint inference of relatedness and the probability of allele sharing for nIBD parasites [41, 56, 57, 58]. We envision an iterative construction — as articulated by Thomas and Hill [56], Smith, Herbinger, and Merry [57], and Wang [58] in the context of sibship reconstruction — whereby relatedness estimates would inform the readjustment of nIBD-to-observation model across successive iterations.

## 5 Conclusion

Based on our results practitioners have two options: resolve to use relative relatedness estimated under independence or try to estimate absolute relatedness under a HMM. Because relative estimates are intrinsically adjusted, caution is required when differences across transmission settings are sought after. That said, if relatedness estimates are viewed as relationship-indicators, relative values are arguably preferable across transmission settings. Caution should be exercised when estimating absolute relatedness because the extent to which underestimation is mitigated is unknowable, but analyses dependent on absolute values and high relatedness thresholds are relatively robust. We are reluctant to prescribe decision thresholds given most use cases are likely to deviate from any contrived examples. Instead, we provide a framework to help practitioners evaluate their individual situations on a case-by-case basis.

## Data availability statement

Sequencing data for *P. falciparum* isolates from Guyana [44] fall under BioProject PRJNA809659; accession numbers are forthcoming. Sequencing data for *P. falciparum* isolates from Colombia [14] are available in the NCBI Sequencing Read Archive; accession numbers are provided in supplemental Table S1. A minimum analysis dataset for the present project, comprising a genotype matrix, is available on the accompanying GitHub repository: https://github.com/somyamehra/PlasmodiumRelatednessBias

## Acknowledgements

We thank our collaborators Dr Reza Niles-Robin, Dr Horace Cox and Dr Collete Clementson from the Ministry of Health, Guyana; Dr Vladimir Corredor from Universidad Nacional de Colombia; and Dr Julian C. Rayner from University of Cambridge. We thank all study participants and technicians who contributed to the data analysed in this project.

DEN is supported by the Bill & Melinda Gates Foundation (investment ID INV-009416). Under the grant conditions of the Foundation, a Creative Commons Attribution 4.0 Generic License has already been assigned to the Author Accepted Manuscript version that might arise from this submission. DEN is also supported with federal funds from the National Institute of Allergy and Infectious Diseases, National Institutes of Health, Department of Health and Human Services, under Grant Number U19AI110818 to the Broad Institute.

MW is supported by the French Government’s Laboratoire d’Excellence “Integrative Biology of Emerging Infectious Diseases” (Investissement d’Avenir grant n°ANR-10-LABX-62-IBEID), and INCEPTION programs (Investissement d’Avenir grant ANR-16-CONV-0005). MW and ART are supported by the P. vivax Serology for Elimination Partnership (Investment ID INV-024368) funded by The Bill & Melinda Gates Foundation.

ART is now funded by the European Union (project number 101110393). Views and opinions expressed are however those of the author only and do not necessarily reflect those of the European Union or European Research Executive Agency (REA). Neither the European Union nor the granting authority can be held responsible for them.

SM gratefully acknowledges a Fulbright Future Scholarship (funded by The Kinghorn Foundation), and the Australian-American Fulbright Commission.

## Notes

https://github.com/somyamehra/PlasmodiumRelatednessBias

### Competing Interest Statement

The authors have declared no competing interest.

