## Appendices A to C for "Systematic bias in malaria parasite relatedness estimation"

**Summary**

This supplement provides a comprehensive account of our theoretical, numerical and empirical analyses. Details of the full study, including intermediate results, are described in Appendix A. The construction of the simulation model is discussed in Appendix B. A glossary of terms used throughout the study is provided in Appendix C.

**C Glossary of terms** **48**

### Appendix A

### Detailed methods and results

| | THEORY | NUMERICAL | EMPIRICAL |
|---|---|---|---|
| **Sample allele frequencies partially encode relatedness structure**<br>*Misspecification of standard (n)IBD-to-observation models* | ✓ | | |
| **Relatedness is systematically underestimated under the independence model**<br>*Re-interpretation as a relative measure, intrinsically adjusted for relatedness averaged over the parasite sample; zero-inflation* | ✓ | ✓ | |
| **Exploiting linkage structure using the HMM of relatedness can mitigate understimation for dense datasets**<br>*Reduced sensitivity to misspecified (n)IBD-to-observation models* | | ✓ | |
| **A dense data diagnostic: gauging the severity of underestimation**<br>*Comparison of dense data estimates under the independence model vs HMM can elucidate average population-level relatedness* | | ✓ | ✓ |
| **Analysing sparse data under the HMM yields an intermediary regime**<br>*Partial but incomplete mitigation of underestimation; may be sufficient to identify highly-related parasite pairs* | | | ✓ |
| **Diagnostics for population structure**<br>*Multimodal empirical IBS distributions and systematic differences in relatedness estimates with IBS descriptives vs allelic states indicate population structure* | | | ✓ |

**Table A.1:** Overview of results, with theoretic, numerical and/or empirical support.

| Quantity | Interpretation | Equation |
|---|---|---|
| $A_i^{(k)}$ | Allelic state for individual $k$ at locus $i$ (observable) | – |
| $S_i^{(k,\ell)}$ | IBS state for individuals $k, \ell$ at locus $i$ (observable) | (A.2) |
| $D_i^{(k,\ell)}$ | IBD state for individuals $k, \ell$ at locus $i$ (unobservable) | – |
| $f_i(q)$ | Sample frequency of allele $q$ at locus $i$ (observable) | (A.1) |
| $d_i$ | Sample proportion of pairs IBD at locus $i$ (unobservable) | (A.8) |
| $s_i$ | Sample proportion of pairs IBS at locus $i$ (observable) | (A.12) |
| $c_i$ | Sample proportion of pairs IBC at locus $i$ (unobservable) | (A.13) |
| $r^{(k,\ell)}$ | Pairwise relatedness parameter for parasites $k$, $\ell$ | (A.19) |
| $\hat{r}^{(k,\ell)}$ | MLE of pairwise relatedness parameter for parasites $k$, $\ell$ | – |

**Table A.2:** Summary of notation and key quantities for Appendix A only. The default for comparative variables includes self-self comparisons (that is, entails sampling with replacement).

#### A.1 Methods

Our approach for characterising systematic biases in malaria parasite relatedness estimation is three-fold. We begin by deriving results under a theoretical framework, geared towards the pairwise estimation of relatedness. We then construct a simulation model designed to capture successive generations of inbreeding, under which we verify our theoretical results and design practical diagnostics. We conclude with a case study of *P. falciparum* data from an inbred parasite population to demonstrate the implications of our findings. Throughout, individual is used to refer to a parasite genotype drawn from an infected host, whereas sample is used to refer to a collection of $k = 1, \ldots, n$ parasite genotypes drawn from many infected hosts. See Table A.2 for a complete list of notation used throughout this appendix. For brevity and clarity of exposition, this notation differs from the main text: specifically, the sample proportion of pairs that are IBD (IBS) at locus $i$ is denoted $d_i$ ($s_i$) here, compared to $\overline{\text{IBD}}_i$ ($\overline{\text{IBS}}_i$) in the main text.

##### A.1.1 Theoretical framework

While allelic concordance or identity-by-state (IBS) is observable, it can be attributed to one of two latent states: identity-by-descent (IBD), reflecting relatedness, i.e., a common ancestral origin; identity-by-chance (IBC), otherwise. Hereafter, we use nIBD as shorthand for 'not IBD'. Note that IBC $\neq$ nIBD; rather, IBC = IBS $\cap$ nIBD. We use (n)IBD as shorthand for both IBD and nIBD, likewise for (n)IBS.

Before proceeding, let us introduce some notation. Given a sample of $n$ not necessarily distinct parasite genotypes, we index each genotype $k = 1, \ldots, n$. We allow each locus $i = 1, \ldots, m$ to harbour an allele in the set $\{1, \ldots, y\}$, where $y$ denotes the maximal cardinality. We denote by $A_i^{(k)}$ the allele observed at locus $i$ in genotype $k$. The sample frequency of allele $q$ at locus $i$ is then calculated to be

$$f_i(q) = \frac{1}{n} \sum_{k=1}^{n} \mathbb{1}\{A_i^{(k)} = q\}, \tag{A.1}$$

where $\mathbb{1}\{\cdot\}$ denotes the indicator function. For a given parasite pair $(k, \ell)$, we further define

- $S_i^{(k,\ell)} = 1$ if locus $i$ is IBS for the pair $(k, \ell)$ and zero otherwise, or equivalently

$$S_i^{(k,\ell)} = \mathbb{1}\{A_i^{(k)} = A_i^{(\ell)}\} = \sum_{q=1}^{y} \mathbb{1}\{A_i^{(k)} = q\}\mathbb{1}\{A_i^{(\ell)} = q\}. \tag{A.2}$$

- $D_i^{(k,\ell)} = 1$ if locus $i$ is IBD for the pair $(k, \ell)$ and zero otherwise.

Throughout, we perform all pairwise comparisons with replacement, i.e, including self-self comparisons.

Relatedness structure within a parasite population is governed by some unknown ancestral stochastic process, replete with demographic complexity. The characterisation and analysis of this ancestral process is beyond our scope. Instead, we seek to analyse a sample of parasite genotypes at a given point in time using a non-ancestral model. In particular, given the set of observed sequences of allelic states $\{\boldsymbol{A}^{(k)}\}_{k=1}^{n}$ for a sample of $n$ individuals, we seek to probabilistically recapitulate the unobservable relatedness structure $\{\boldsymbol{D}^{(k,\ell)}\}_{k=1,\ell<k}^{n}$ (Figure A.1).

###### A.1.1.1 Joint model of malaria parasite relatedness

In an ideal setting, we would perform joint inference over the sample of $n$ individuals [10, 26]. The construction of a joint model, however, is highly non-trivial. The principal complication lies in the non-independence of IBD states $\boldsymbol{D}^{(k,\ell)}$ between pairs of individuals. Since IBD is a transitive property, rather than considering parasite pairs in isolation, we would need to construct graphs of parasite relationships on the population-level [27]. In reconstructing the complete relatedness structure of a population genotyped at $m$ loci, we would recover a system of $m$ correlated, transitive graphs – each representing IBD sharing at a given locus – with the number of allowable configurations per graph growing as Bell numbers as a function of the parasite population size. This poses a significant combinatorial problem, quickly rendering brute force approaches computationally intractable [27].

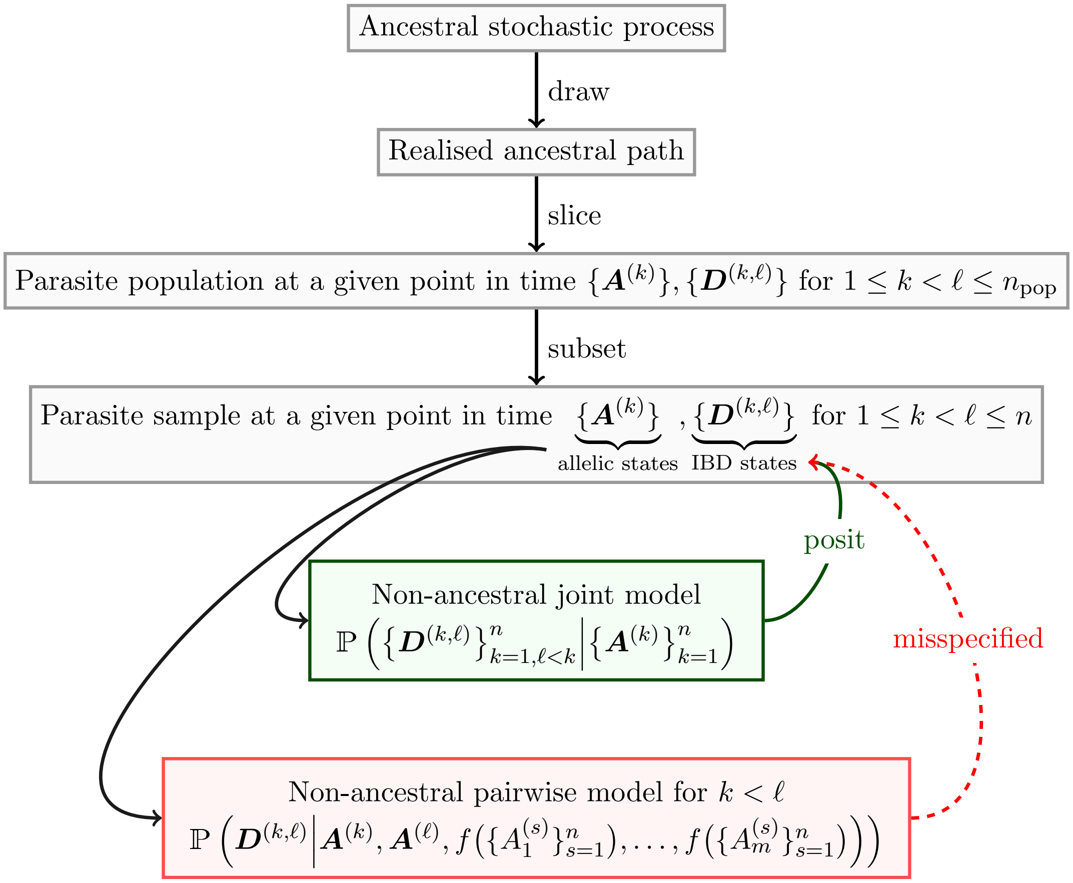

**Figure A.1:** Conceptual overview, where $f\left(\{A_i^{(s)}\}_{s=1}^{n}\right)$ denotes the vector of sample allele frequencies for alleles $\{1, \ldots, y\}$ at locus $i$.

###### A.1.1.2 Non-ancestral pairwise Markov model of (n)IBD states

In practice, relatedness is estimated using a non-ancestral pairwise model, comprising an alternating Poisson process (APP) [3]. For a given parasite pair, we conceptualise the genome as a mosaic of alternating (n)IBD segments, with the intution that IBD tracts are fragmented by randomly-distributed recombination breakpoints over successive generations [3].

While the genome is treated to be continuous under the APP model, we sample a finite number of discrete loci $i = 1, \ldots, m$. To describe the sequence of IBD states

$$\boldsymbol{D} = (D_1, \ldots, D_m) \in \{0, 1\}^m$$

across loci $i = 1, \ldots, m$ (where parasite identifiers $k, \ell$ are dropped for notational convenience), Leutenegger et al. [9] construct a discrete-time Markov process. Linkage between successive markers is parametrised by:

- the genomic distance $\delta_i$ in units of base pairs (bp) between loci $i$ and $(i+1)$;
- a constant recombination rate $\rho$ in units of Morgans per base pair (M/bp), typically ascertained from genetic cross experiments [16, 18]; and
- a (n)IBD latent state switching rate $\kappa$ that is unobservable and must be inferred.

We assume that locus 1 is IBD with probability $r$, that is,

$$\mathbb{P}(D_1 = 1) = r,\ \mathbb{P}(D_1 = 0) = 1 - r.$$

Under the Markov property, $D_{i+1}$ is dependent only on $D_i$. The complete sequence of IBD states $\boldsymbol{D}$ is governed by the transition matrix

$$\begin{pmatrix} \mathbb{P}\left(D_{i+1}(t) = 0 | D_i(t) = 0\right) & \mathbb{P}\left(D_{i+1}(t) = 1 | D_i(t) = 0\right) \\ \mathbb{P}\left(D_{i+1}(t) = 0 | D_i(t) = 1\right) & \mathbb{P}\left(D_{i+1}(t) = 1 | D_i(t) = 1\right) \end{pmatrix}$$
$$:= \begin{pmatrix} 1 - r(1 - e^{-\kappa\rho\delta_i}) & r(1 - e^{-\kappa\rho\delta_i}) \\ (1-r)(1 - e^{-\kappa\rho\delta_i}) & 1 - (1-r)(1 - e^{-\kappa\rho\delta_i}) \end{pmatrix}, \tag{A.3}$$

as per Taylor et al. [26], where $k$ was used instead of $\kappa$ for the switch rate parameter. The degree of dependence between successive loci is governed by the product $\kappa\rho\delta_i$: to weaken the dependence between successive loci, we can either increase the switching rate $\kappa$, or the genomic distance $\delta_i$ between successive loci.

###### A.1.1.3 Non-ancestral pairwise independence model of (n)IBD states

Under the above-mentioned non-ancestral Markov model in the limit $\kappa \to \infty$ [26] or $\delta_i \to \infty$ [20], we recover the independence model

$$\mathbb{P}(\boldsymbol{D}) = \prod_{i=1}^{m} r^{D_i}(1-r)^{1-D_i}, \tag{A.4}$$

where (n)IBD states are described by independent and identically distributed Bernoulli random variables with success probability $r$.

###### A.1.1.4 Observation models

Since (n)IBD states are unobservable, estimation of the pairwise relatedness parameter $r$ necessitates coupling the model of hidden (n)IBD states to a model of observations conditional on (n)IBD states. Herein, observations are either alleles or (n)IBS descriptives (Box 1). We do not model genotyping errors. Elsewhere, where genotyping errors are taken into account (e.g., Taylor et al. [26]), the observation model integrates over latent non-erroneous alleles and is thus made of modules: an (n)IBD-to-latent-allele observation model, and a model capturing genotyping error, e.g.,

$$\mathbb{P}\left(\boldsymbol{A}^{(k)}_{\text{obs}}, \boldsymbol{A}^{(\ell)}_{\text{obs}} \middle| \boldsymbol{D}^{(k,\ell)}\right) = \sum_{\boldsymbol{A}^{(k)},\boldsymbol{A}^{(\ell)}} \underbrace{\mathbb{P}\left(\boldsymbol{A}^{(k)}, \boldsymbol{A}^{(\ell)} \middle| \boldsymbol{D}^{(k,\ell)}\right)}_{\substack{\text{(n)IBD-to-latent-allele}\\ \text{model}}} \times \underbrace{\mathbb{P}\left(\boldsymbol{A}^{(k)}_{\text{obs}}, \boldsymbol{A}^{(\ell)}_{\text{obs}} \middle| \boldsymbol{A}^{(k)}, \boldsymbol{A}^{(\ell)}\right)}_{\text{error model}}.$$

Further details of the observations models are provided in the results section. They include standard practice observation models (models into which sample allele frequencies are plugged, where sample allele frequencies may be computed before or after removing replicates of seemingly clonal parasites); and a corrected model of independent (n)IBS states, which is not practically available, but useful for validating theory when applied to simulated data.

**Box 1: Observations are either alleles or (n)IBS descriptives**

Consider the following data on a pair of individual genotypes

*genotype 1*: $A$ $T$ $C$ $G$

*genotype 2*: $A$ $C$ $C$ $G$

The observations on the pair of genotypes 1 and 2 are

| | | | | |
|---|---|---|---|---|
| $(A, A)$ | $(T, C)$ | $(C, C)$ | $(G, G)$ | using alleles |
| IBS | nIBS | IBS | IBS | using (n)IBS descriptives |

##### A.1.2 Simulated data

The model under which simulated data are generated is described in detail in Appendix B. Assumed parameter values, and their respective interpretations, are detailed in Table B.1. The model's purpose is not to recapitulate epidemiological reality, but to generate data that can be used to verify theoretical results and facilitate the design of practical diagnostics. To ensure verification is independent, data are simulated under a model built on ancestral principles; they are not simulated under the non-ancestral models used to estimate relatedness.

Under the simulation model, there is a dichotomy between ancient low-level background relatedness stemming from generation zero, and very recent relatedness arising from intensive inbreeding under a small fixed population size. To balance the distribution of IBD segment lengths in spite of this dichotomy, we use a recombination rate and genome size that substantially exceeds that of *P. falciparum* [18].

Estimates of pairwise relatedness generated under the non-ancestral relatedness models are compared with true pairwise relatedness values, which are known for simulated data. For a given pair of simulated individuals, we use an approximation of realised relatedness as our truth value, where realised relatedness is the fraction of loci that are IBD [17]. Under our simulation model where markers are equidistant, realised relatedness is approximated by the fraction of polymorphic markers that are IBD: IBD is ascertained by comparing ancestral founder mosaics for each pair of simulated individuals; polymorphic markers are those at which there is some variation within the sample, noting that the number of polymorphic markers for a given sample tends to decrease over generations due to loss of diversity. We compute realised relatedness using polymorphic markers because pairwise relatedness estimates are predicated on data from polymorphic markers only.

***Aside***: In light of the finite length of the genome, there is a fundamental difference between a pairwise relatedness parameter $r$ and realised relatedness [17, 26], although we expect convergence of realised relatedness to $r$ for an infinite number of equidistant markers along an infinitely-long genome. However, since we simulate recombination from ancestral principles, rather than simulating IBD states under the non-ancestral HMM of relatedness or its independent counterpart, there is no truth parameter equivalent to $r$ under our simulation framework.

##### A.1.3 *P. falciparum* data

We illustrate the practical consequences of our theoretical and numerical findings through a case study of a highly-inbred parasite population. We focus on a set of highly-quality isolates from passively sampled symptomatic patients in Guyana in 2016–2020 [36], as well as from Colombia in 1993–2017 [33], that are deemed to be monoclonal. WGS data for these isolates were processed previously in accordance with GATK best practices to yield a genomewide set of variants as described in Carrasquilla et al. [33] and Vanhove et al. [36]. Given the complete set of WGS variants, we apply additional filtration criteria:

- Indels and multiallelic variants are removed to retain only biallelic SNPs (i.e., with precisely one alternate allele) that are polymorphic among the sampled infections.
- SNPs lying in centromeric, hypervariable or subtelomeric regions of the *P. falciparum genome* (as defined by Miles et al. [18]) are removed, yielding variants in the core nuclear genome only.
- Individual variant calls that are either heterozygous or have read support below 5 (based on the DP tag) are masked (i.e. set to be missing).
- Variant sites with missingness $> 30\%$ across isolates are removed.
- Isolates with missingness $> 30\%$ across filtered variant sites are removed.

This yields a WGS SNP dataset comprising $n = 306$ isolates ($n = 278$ from Guyana, $n = 28$ from Colombia) and $n = 30694$ polymorphic biallelic SNPs. Using the dataset, we elucidate the consequences of population structure, which has been omitted from our theoretical and numerical analyses. We also illustrate the implications of marker sparsity on pairwise relatedness estimates for $n = 278$ isolates from Guyana (genotyped at $n = 16115$ polymorphic biallelic SNPs), generating sparse marker panels by down-sampling SNPs sites uniformly at random without replacement.

#### A.2 Theoretical results

Standard (n)IBD-to-observation models are constructed using sample allele frequencies. When relatedness structure is significant, overlooking it in allele frequency estimation can introduce significant biases [7, 8, 10, 11, 14, 21], with practical consequences including the systematic underestimation of relatedness [13] (Case A, Box A.2). Removing replicates of seemingly clonal parasites from the sample before computing allele frequencies re-weights allele frequencies in a case-specific manner. It does not account for relatedness between the remaining sample members, however. As such, systematic underestimation of relatedness is liable to persist (Case A, Box 2). An alternative naive approach, evoking the assumption of equifrequent alleles, may yield systematic overestimates of pairwise relatedness [13] (Case B, Box A.2).

**Box 2: Some consequences of misspecified observation models**

**Likelihood of pairwise IBS sharing for a given locus assuming no error**

$$\begin{aligned}\mathbb{P}(\mathrm{IBS}) &= \mathbb{P}(\mathrm{IBS}\,|\,\mathrm{IBD})\,\mathbb{P}(\mathrm{IBD}) + \mathbb{P}(\mathrm{IBS}\,|\,\mathrm{nIBD})\,\mathbb{P}(\mathrm{nIBD}) \\ &= \mathbb{P}(\mathrm{IBD}) + \mathbb{P}(\mathrm{IBS}\,|\,\mathrm{nIBD})\Big(1 - \mathbb{P}(\mathrm{IBD})\Big)\end{aligned}$$

since $\mathbb{P}(\mathrm{IBS}\,|\,\mathrm{IBD}) = 1$ (in the absence of genotyping error), and where $\mathbb{P}(\mathrm{IBD})$ is a genomewide measure of relatedness.

**Implications for relatedness estimates**

- $\mathbb{P}(\mathrm{IBS}\,|\,\mathrm{nIBD})$ overestimated $\implies$ $\mathbb{P}(\mathrm{IBD})$ underestimated
- $\mathbb{P}(\mathrm{IBS}\,|\,\mathrm{nIBD})$ underestimated $\implies$ $\mathbb{P}(\mathrm{IBD})$ overestimated

**Observation models**

(A) $\mathbb{P}(\mathrm{IBS}\,|\,\mathrm{nIBD}) \approx \overline{\mathrm{IBS}}$, the observed proportion of IBS pairs in the sample*

| | |
|---|---|
| *Calculation:* | sums of squares of sample allele frequencies |
| *Possible issue:* | IBS due to IBD is not attributed to IBD |
| *Implication:* | $\mathbb{P}(\text{IBS} \mid \text{nIBD})$ may be overestimated |

(B) $\mathbb{P}(\text{IBS} \mid \text{nIBD}) \approx 1/n$ where $n$ is locus cardinality

| | |
|---|---|
| *Calculation:* | reciprocal of observed allele count |
| *Possible issue:* | alleles are not equifrequent |
| *Implication:* | $\mathbb{P}(\text{IBS} \mid \text{nIBD})$ may be underestimated [2] |

* replicates of clonal parasites may have been removed from the sample or not.

Starting with the standard (n)IBD-to-allele model, and its corrected IBD-to-allele counterpart (Section A.2.1), we show that standard observation models can be misspecified given both IBD (Section A.2.1.1) and nIBD (Sections A.2.1.2, A.2.2.1, A.2.2.2), and reason why removing replicates of seemingly clonal parasites before computing allele frequencies rectifies neither case. More specifically our theoretical results show that, given IBD, standard observation models are potentially misspecified due to assumed independence between IBD and allelic states, which does not always hold (Section A.2.1.1); given nIBD they are misspecified due to partial encoding of the locuswise proportion of IBD pairs within the sums of squares of sample allele frequencies [21] (Section A.2.1.2). Systematic underestimation of pairwise relatedness may be associated with this misspecification. Under the independence model of relatedness with IBS descriptives, we explore additional consequences of this misspecification (Section A.2.2). Firstly, echoing the work of Weir and Goudet [21] (who also adopt IBS descriptives), we re-interpret the pairwise relatedness parameter $r$ as a relative measure, capturing deviation from average relatedness (Section A.2.2.1). Secondly, we show pairwise relatedness estimates are stratified by average relatedness, with zero-valued estimates below and positive estimates above (Section A.2.2.2), likewise echoing Weir and Goudet [21]. While our theoretical results are restricted to the independence model, the implications of marker density and linkage are explored through analyses of simulated and empirical data (Sections A.3 and A.4, respectively).

##### A.2.1 The standard (n)IBD-to-allele model and its corrected IBD-to-allele counterpart

The construction of the standard (n)IBD-to-allele model [22, 23, 26, 29] is two-fold:

1. Given a pair of parasites $(k, \ell)$ is IBD at locus $i$, the probability that both individuals have

allele $q$ is the sample frequency of allele $q$ at locus $i$:

$$\mathbb{P}_{\text{standard}}\left(A_i^{(k)} = A_i^{(\ell)} = q \,\middle|\, D_i^{(k,\ell)} = 1\right) = f_i(q). \tag{A.5}$$

2. Given a pair of parasites $(k, \ell)$ is nIBD at locus $i$, the probability of observing alleles $(q, u)$ is the product of the sample frequencies of alleles $q$ and $u$ at locus $i$:

$$\mathbb{P}_{\text{standard}}\left(A_i^{(k)} = q, A_i^{(k)} = u \,\middle|\, D_i^{(k,\ell)} = 0\right) = f_i(q) \cdot f_i(u), \tag{A.6}$$

where $f_i$ is a function of $1, \ldots, n$ individuals (Equation (A.1)).

In other words, it is assumed that allele frequencies, when restricted to IBD pairs, are equivalent to sample allele frequencies (Equation (A.5)); and the likelihood of each pairwise allelic state, when restricted to non-IBD pairs, is equivalent to the sample incidence of that pairwise allelic state (Equation (A.6)).

This description applies when allele frequencies are computed using samples from which replicates of seemingly clonal parasites have been removed or not. As such, all results that follow apply to either case. The only difference between the cases is the specific values of $f_i(q)$, $s_i$, $d_i$ etc., which will differ on a case-by-case basis not amenable to generalisation.

To highlight the misspecification of standard observation models, it is instructive to introduce some unobservable quantities of interest:

- The sample proportion of pairs IBD at locus $i$ harbouring allele $q$

$$d_i(q) = \frac{1}{n^2} \sum_{k=1}^{n} \sum_{\ell=1}^{n} \mathbb{1}\{A_i^{(\ell)} = q\} \cdot D_i^{(k,\ell)} \tag{A.7}$$

  in which the indicator features only once since $D_i^{(k,\ell)} = 1$ implies $A_i^{(k)} = A_i^{(l)}$.

- The sample proportion of pairs IBD at locus $i$, i.e., the locuswise average relatedness

$$d_i = \sum_{q=1}^{y} d_i(q) = \frac{1}{n^2} \sum_{k=1}^{n} \sum_{\ell=1}^{n} D_i^{(k,\ell)}. \tag{A.8}$$

The sample proportion of individuals that are IBD at locus $i$ and harbour allele $q$ is $d_i(q)/d_i$. If it were observable, $d_i(q)/d_i$ would be used to approximate the true probability of an IBD pair harbouring allele $q$ under an IBD-to-allele model, since we would expect the sample proportion to converge to the true probability as the sample size $n$ goes to infinity, at least under the conventional law of large numbers. As such, we claim that

$$\mathbb{P}_{\text{corrected}}\left(A_i^{(k)} = A_i^{(\ell)} = q \,\middle|\, D_i^{(k,\ell)} = 1\right) \approx \frac{d_i(q)}{d_i}. \tag{A.9}$$

###### A.2.1.1 The standard IBD-to-allele model assumes independence between IBD and allelic states

When IBD and allelic states are independent, $d_i(q) = f_i(q) \cdot d_i$, rendering

$$\mathbb{P}_{\text{corrected}}\left(A_i^{(k)} = A_i^{(\ell)} = q \mid D_i^{(k,\ell)} = 1\right) \approx f_i(q). \tag{A.10}$$

Otherwise stated, the standard IBD-to-allele model (Equation (A.5)) assumes independence between allelic and IBD states. If independence between allelic and IBD states does not hold, then the sample allele frequency $f_i(q)$ is not necessarily a good approximation of the true probability of an IBD pair harbouring allele $q$, and the standard IBD-to-allele model is misspecified.

The presence of relatedness structure within the sample can violate the assumption of independence between allelic and IBD states. By relatedness structure, we refer to the locuswise partitioning of individuals into IBD clusters [27]. If sub-cluster sizes are equal, then Equation (A.5) is guaranteed to hold. Unbalanced sub-cluster sizes, however, can lead to substantial divergence between the sample proportion of IBD pairs harbouring each allelic state and the likelihood under the standard IBD-to-allele model (Equation (A.5)) (see Figure A.2 for an example); this may occur if a particular allele is under strong positive selection. In the context of sibship reconstruction, which likewise involves partitioning samples by ancestry, the correction of sample allele frequencies is posited to be particularly pertinent in small populations with variable family/partition sizes [5, 6]; an analogous prescription applies here. Unlike family size, relatedness structure within a parasite population is unobservable and poses significant combinatorial challenges for inference: given dependence between parasite pairs, inference of relatedness and allele frequencies should be performed jointly over parasite population. Without due consideration of underlying demographic processes and an associated mechanistic model of ancestry, it is difficult to gauge the expected relatedness structure within a population. Diagnosing the degree of misspecification under the model likelihood (Equation (A.5)) is therefore non-trivial in practice.

Besides in the unrealistic scenario where the parasite population is split into unrelated clonal clusters, removing replicates of seemingly clonal parasites will not remove replicates within locus-wise IBD partitions because individuals can be IBD at a given locus without being IBD at all loci (Figure A.3). Otherwise stated, removing replicates of seemingly clonal parasites from a sample does not remove relatedness between the remaining sample members.

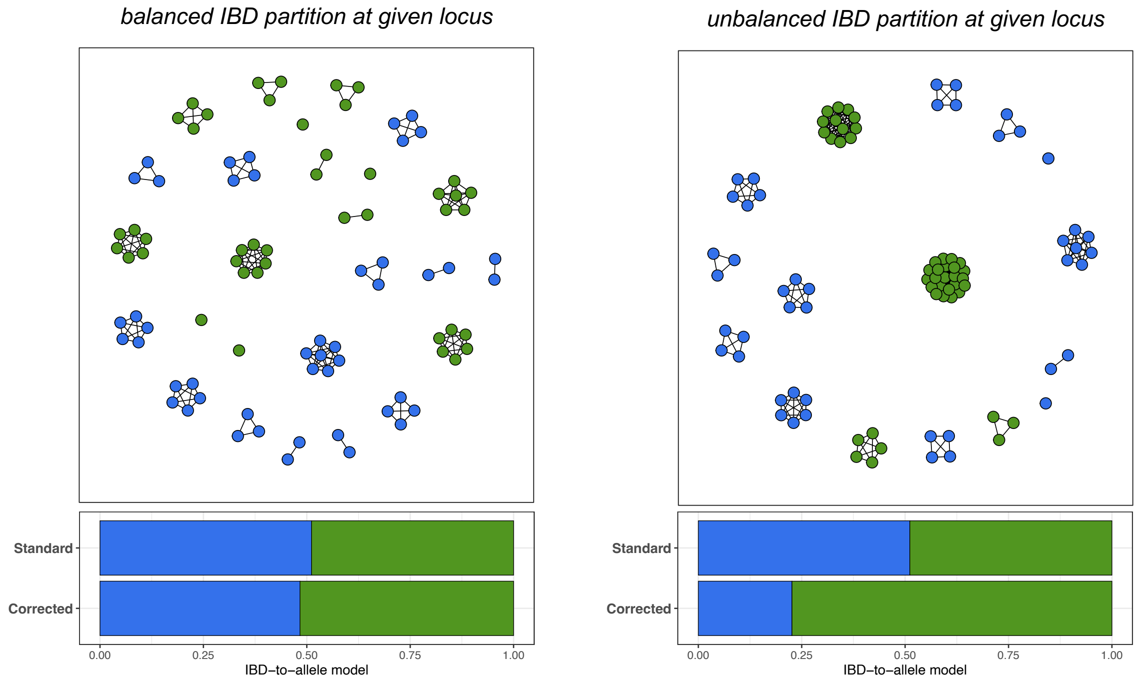

**Figure A.2:** Schematic of misspecification under the standard IBD-to-allele model, relative to the proposed correction (A.9), based on an underlying IBD network for a single locus. Each node represents an individual, coloured by allelic state. Individuals that are IBD at that locus are connected by an edge. Since IBD is a transitive property [27], all components are fully-connected. In the absence of genotyping error, edges can only be placed between nodes of the same colour (that is, IBD necessarily implies IBS).

###### A.2.1.2 The standard nIBD-to-IBS model encodes locuswise average relatedness

The observation model of allelic states for nIBD loci (Equation (A.6)), is misspecified in the conflation of IBC and IBS: an unobservable and unaccounted proportion of IBS is driven by IBD. Similarly to Weir and Goudet [21], we shift to IBS descriptives hereafter to:

1. Circumvent misspecification arising from the assumed independence of allelic and IBD states implicit in Equation (A.5); and
2. Highlight the confounding effects of relatedness structure in the construction of observation models for nIBD pairs (Equation (A.6)).

Condensing Equation (A.6) to consider only IBS descriptives yields the following nIBD-to-IBS model:

$$\mathbb{P}_{\text{standard}}\left(S_i^{(k,\ell)} = 1 \,\middle|\, D_i^{(k,\ell)} = 0\right) = \sum_{q=1}^{y} f_i(q)^2 =: s_i. \tag{A.11}$$

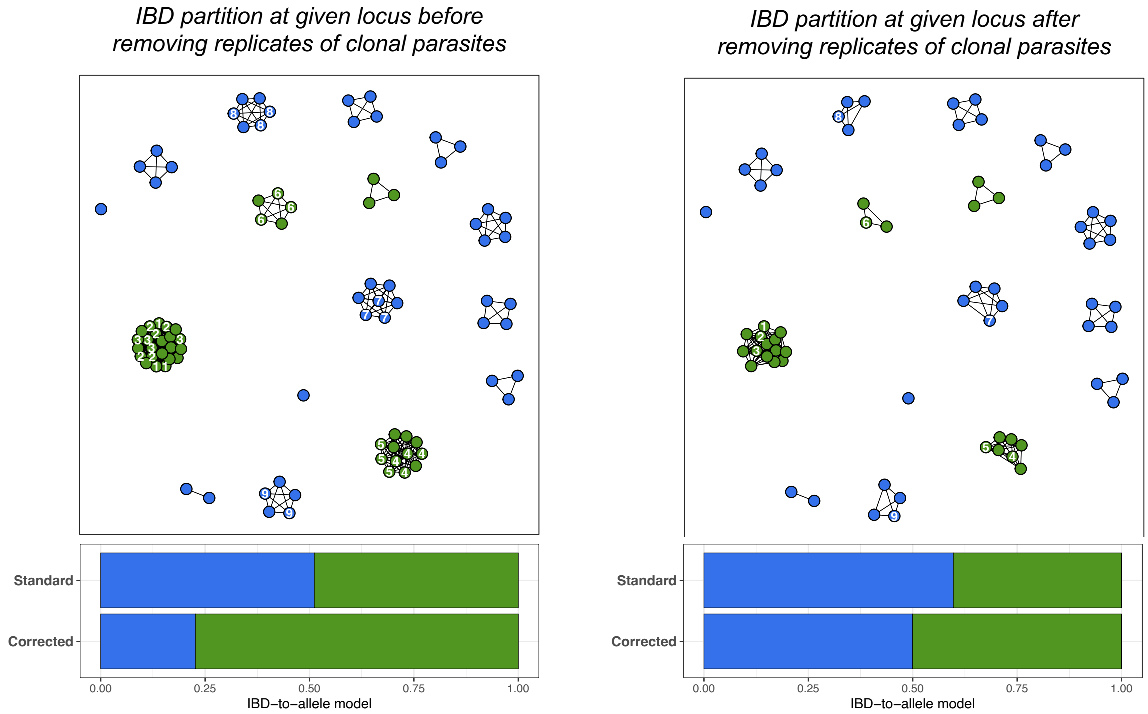

**Figure A.3:** Unbalanced IBD partitions at a given locus before and after removing replicates of clonal parasites. Groups of clonal parasites are numbered 1 to 9.

We note that Equation (A.11) is precisely Nei's gene identity metric [1, 26]. We can alternatively interpret $s_i$ as an estimate of the probability that two identical alleles are selected from a pool of $n$ alleles $A_i^{(k)}$, $k = 1, \dots, n$ *with* replacement by rearranging Equation (A.11) to obtain the expression

$$s_i := \sum_{q=1}^{y} \left( \frac{1}{n} \sum_{k=1}^{n} \mathbb{1}\{A_i^{(k)} = q\} \right)^2 = \frac{1}{n^2} \sum_{k=1}^{n} \sum_{\ell=1}^{n} \left( \sum_{q=1}^{y} \mathbb{1}\{A_i^{(k)} = A_i^{(\ell)} = q\} \right) = \frac{1}{n^2} \sum_{k=1}^{n} \sum_{\ell=1}^{n} S_i^{(k,\ell)}, \tag{A.12}$$

where we have substituted Equations (A.1) and (A.2) and interchanged the order of summation and products.

In the absence of genotyping error, such that locus $i$ can only be IBD if it is IBS, that is, $S_i^{(k,\ell)} \geq D_i^{(k,\ell)}$, the sample proportion of nIBD individuals that are IBS at locus $i$ can be written

$$c_i := \frac{\sum_{k=1}^{n} \sum_{\ell=1}^{n} \left[ S_i^{(k,\ell)} - D_i^{(k,\ell)} \right]}{\sum_{k=1}^{n} \sum_{\ell=1}^{n} \left[ 1 - D_i^{(k,\ell)} \right]} = \frac{s_i - d_i}{1 - d_i}, \tag{A.13}$$

where we have used Equations (A.8) and (A.12).

If $c_i$ were observable, the probability of IBS sharing under an nIBD-to-IBS model would be approximated by it, with analogous reasoning to Section A.2.1.1: in the infinite sample size limit, that is, as $n$ approaches infinity, $c_i$ would be expected to converge to the true conditional probability of IBS given nIBD (at least in the context of the standard law of large numbers). As such, we purport

$$\mathbb{P}_{\text{corrected}}\left(S_i^{(k,\ell)} = 1 \,|\, D_i^{(k,\ell)} = 0\right) \approx c_i. \tag{A.14}$$

This construction allows us to capitulate the misspecification inherent in the standard nIBD-to-IBS model. In particular, rearranging Equation (A.13), we see that the sample proportion of IBS pairs $s_i$ — the standard nIBD-to-IBS model likelihood — can be written as a function of the unobservable proportion of IBD pairs $d_i$ and the unobservable sample proportion of nIBD individuals that are IBS $c_i$ as follows:

$$\text{Standard nIBD-to-IBS model} = s_i = c_i + (1 - c_i)d_i \neq c_i. \tag{A.15}$$

Otherwise stated, $d_i$ partially encodes relatedness structure in the sample proportion of IBS pairs $s_i$ and thus the standard nIBD-to-IBS model.

Since pairwise comparisons are performed with replacement, i.e, self-self comparions are included, it is necessarily the case that $d_i \geq 1/n$, whereby $s_i \geq c_i$. As such, the standard nIBD-to-IBS model overestimates the probability of IBS for nIBD pairs at locus $i$. The degree of overestimation grows linearly as a function of the proportion of pairs $d_i$ that are IBD at locus $i$ – and may therefore be particularly problematic in populations with high levels of relatedness.

***Aside***: Purcell et al. [12] propose a correction for finite sample bias that is predicated on the probability that two identical alleles are selected from a pool of $n$ alleles *without* replacement, i.e, excluding self-self comparisons. Implementing this correction yields the adjusted quantity

$$\text{Adjusted nIBD-to-IBS model} = s_i' := \frac{1}{\binom{n}{2}} \sum_{k=1}^{n-1} \sum_{\ell=k+1}^{n} S_i^{(k,\ell)} = \frac{s_i - \frac{1}{n}}{1 - \frac{1}{n}}, \tag{A.16}$$

where we have substituted Equation (A.12). For sufficiently large sample sizes $n$, both formulations $s_i \approx s_i'$ are approximately equal. This adjustment is sufficient in an outbred setting where all pairs of distinct parasites are unrelated, but falls short for inbred parasite samples.

***Aside***: While we have adopted the convention of pairwise comparisons with replacement, in line with standard nIBD-to-observation models [22, 23, 26], this decision does not affect the theoretical correction $c_i$. That is, if we define

$$d_i' := \frac{1}{\binom{n}{2}} \sum_{k=1}^{n-1} \sum_{\ell=k+1}^{n} D_i^{(k,\ell)}$$

to be the proportion of distinct pairs that are IBD at locus $i$, then analogous reasoning yields the corrected nIBD-to-IBS model

$$c_i' = \frac{\sum_{k=1}^{n-1} \sum_{\ell=k+1}^{n} \left[S_i^{(k,\ell)} - D_i^{(k,\ell)}\right]}{\sum_{k=1}^{n-1} \sum_{\ell=k+1}^{n} \left[1 - D_i^{(k,\ell)}\right]} = \frac{s_i' - d_i'}{1 - d_i'}. \tag{A.17}$$

Inspection of Equations (A.13) and (A.17), however, reveals that $c_i = c_i'$.

***Aside***: As for misspecification given IBD (Section A.2.1.1), besides in the unrealistic scenario where the parasite population is split into unrelated clonal clusters — in which case the finite-sample correction (A.16) of Purcell et al. [12] is directly applicable — removing replicates of clonal parasite from a parasite sample will not render $d_i$ equal zero, because individuals that are IBD at the $i$th locus are not necessarily IBD at all loci. Otherwise stated, removing replicates of seemingly clonal parasites from a sample does not remove relatedness between the remaining sample members.

##### A.2.2 Consequences of the standard nIBD-to-IBS model under marker independence

The misspecification of the standard nIBD-to-IBS model leads to double-counting relatedness in the likelihood of IBS:

$$\mathbb{P}(\text{IBS}) = \mathbb{P}(\text{IBD}) + \underbrace{\overbrace{\mathbb{P}_{\text{standard}}(\text{IBS}\big|\text{nIBD})}^{\substack{\text{obs model: inflated by} \\ \text{locuswise proportion IBD}}} \cdot \big[1 - \mathbb{P}(\text{IBD})\big]}_{\text{likelihood of IBC} = \text{IBS} \cap \text{nIBD}};$$

intuitively, we expect this to lead to the systematic underestimation of pairwise relatedness (Case A, Box 1). To characterise more thoroughly the consequences of this misspecification, we restrict our attention to the loci-independence model of pairwise relatedness [26] with IBS descriptives. For the parasite pair $(k, \ell)$, the likelihood of observing the sequence of IBS states

$$\mathbf{S}^{(k,\ell)} = \left(S_1^{(k,\ell)}, \dots, S_m^{(k,\ell)}\right)$$

across a set of loci $i = 1, \dots m$, conditional on pairwise relatednes $r^{(k,\ell)}$, is given by

$$\mathbb{P}_{\text{standard}}\left(\boldsymbol{S}^{(k,\ell)} | r^{(k,\ell)}\right) = \prod_{i=1}^{m} \left(r^{(k,\ell)} + s_i(1 - r^{(k,\ell)})\right)^{S_i^{(k,\ell)}} \left(1 - r^{(k,\ell)} - s_i(1 - r^{(k,\ell)})\right)^{1 - S_i^{(k,\ell)}}. \tag{A.18}$$

The results in the present section bear strong conceptual similarity to the work of Weir and Goudet [21]; however, we consider maximum likelihood estimates whilst Weir and Goudet [21] addressed method of moments estimators of relatedness.

###### A.2.2.1 The pairwise relatedness parameter captures deviation from average relatedness

To aid reinterpretation by inspection, we rewrite the likelihood (A.18) in the following form:

$$\begin{aligned}
\mathbb{P}_{\text{standard}}\left(\boldsymbol{S}^{(k,\ell)}|r^{(k,\ell)}\right) &= \prod_{i=1}^{m}\left[1-(1-s_i)(1-r^{(k,\ell)})\right]^{S_i^{(k,\ell)}}\left[(1-s_i)(1-r^{(k,\ell)})\right]^{1-S_i^{(k,\ell)}}, \\
&= \prod_{i=1}^{m}\Big\{\left[1-(1-c_i)\left\{1-d_i-r^{(k,\ell)}(1-d_i)\right\}\right]^{S_i^{(k,\ell)}} \\
&\qquad \left[(1-c_i)\left\{1-d_i-r^{(k,\ell)}(1-d_i)\right\}\right]^{1-S_i^{(k,\ell)}}\Big\},
\end{aligned} \tag{A.19}$$

where $s_i = c_i + (1-c_i)d_i$ (Equation (A.15)) has been substituted into the first line.

Under the corrected nIBD-to-IBS model, $c_i$ (Equation (A.13)), the probability of $\boldsymbol{S}^{(k,\ell)}$ is analogously given by

$$\mathbb{P}_{\text{corrected}}\left(\boldsymbol{S}^{(k,\ell)}|r^{(k,\ell)}_{\text{corrected}}\right) = \prod_{i=1}^{m}\left[1-(1-c_i)(1-r^{(k,\ell)}_{\text{corrected}})\right]^{S_i^{(k,\ell)}}\left[(1-c_i)(1-r^{(k,\ell)}_{\text{corrected}})\right]^{1-S_i^{(k,\ell)}}. \tag{A.20}$$

From inspection of Equations (A.19) and (A.20), the probability of IBD sharing at locus $i$ under the corrected model can be written in the form

$$\mathbb{P}_{\text{corrected}}\left(D_i^{(k,\ell)}=1\right) = \underbrace{d_i}_{\substack{\text{average relatedness}\\ \text{in parasite sample}}} + \underbrace{r^{(k,\ell)}(1-d_i)}_{\substack{\text{deviation from average relatedness}\\ \text{in parasite sample}}}. \tag{A.21}$$

***Aside***: We do not equate $r^{(k,\ell)}_{\text{corrected}}$ to $r^{(k,\ell)}$ because $d_i$ are not necessarily the same for all $i$.

We suggest that Equation (A.21) captures the "true" likelihood of IBD sharing at locus $i$, and can be decomposed into two components:

1. the sample proportion of IBD pairs $d_i$ reflecting the average relatedness at locus $i$ in the parasite sample; and

2. the term $r^{(k,\ell)}(1-d_i)$ quantifying pairwise IBD sharing at locus $i$ that can not be explained by the average locuswise relatedness in the parasite sample alone.

In practice, inference is performed on the parameter $r^{(k,\ell)}$. Rearranging Equation (A.21), we can write

$$r^{(k,\ell)} = \frac{\mathbb{P}_{\text{corrected}}\left(D_i^{(k,\ell)}=1\right)-d_i}{1-d_i}. \tag{A.22}$$

Equation (A.22) suggests that the parameter $r^{(k,\ell)}$ is intrinsically adjusted for locuswise average relatedness in the parasite sample, $d_i$, which varies across loci. In particular, we suggest that $r^{(k,\ell)}$ should be interpreted as a relative measure, quantifying how strongly relatedness between the parasite pair $(k,\ell)$ deviates from the locuswise average relatedness $d_i$ in the parasite sample. This mirrors the notion of relative relatedness articulated by Weir and Goudet [21].

###### A.2.2.2 Relatedness estimates are stratified by average relatedness: zero below, positive otherwise

In practice, the pairwise relatedness parameter $r^{(k,\ell)}$ is taken to have range $[0,1]$. From Equation (A.21), however, we see that the constraint $r^{(k,\ell)} \geq 0$ introduces a lower bound

$$\mathbb{P}_{\text{corrected}}\left(D_i^{(k,\ell)} = 1\right) \geq d_i$$

on the "corrected" model likelihood of IBD sharing at locus $i$ for the parasite pair $(k,\ell)$. In particular, setting $r^{(k,\ell)} = 0$ is equivalent to approximating the locuswise probability of pairwise IBD by the locuswise sample average relatedness $d_i$, whereby

$$\mathbb{P}_{\text{standard}}\left(\mathbf{D}^{(k,\ell)} | r^{(k,\ell)} = 0\right) = \prod_{i=1}^{m} d_i^{D_i}(1-d_i)^{1-D_i}$$

and we recover the expected distribution of pairwise relatedness for the parasite genotype sample. This observation echoes Weir and Goudet [21], who additionally allude to the concept of negative relatedness.

The notion of ostensibly "unrelated" parasite pairs $r^{(k,\ell)} = 0$ thus warrants attention. To explore this notion, it is instructive to consider a MLE scheme for the parameter $r_{k,\ell}$. From Equation (A.19), we can readily verify that the derivative of the log-likelihood for the sequence of observed IBS states $\boldsymbol{S}^{(k,\ell)}$

$$\frac{d \log \mathbb{P}_{\text{standard}}\left(\boldsymbol{S}^{(k,\ell)} | r^{(k,\ell)}\right)}{dr^{(k,\ell)}} = \sum_{i=1}^{m} \left\{ \frac{(1-s_i)S_i^{(k,\ell)}}{s_i + (1-s_i)r^{(k,\ell)}} - \frac{1-S_i^{(k,\ell)}}{1-r^{(k,\ell)}} \right\} \tag{A.23}$$

is a monotonically decreasing function of $r^{(k,\ell)}$. The derivative of the log-likelihood (A.23) allows us to identify a sufficient and necessary condition to recover a zero MLE $\hat{r}^{(k,\ell)} = 0$:

$$\left.\frac{d \log \mathbb{P}_{\text{standard}}\left(\boldsymbol{S}^{(k,\ell)} | r^{(k,\ell)}\right)}{dr^{(k,\ell)}}\right|_{r^{(k,\ell)}=0} \leq 0 \iff \frac{1}{m}\sum_{i=1}^{m} \frac{S_i^{(k,\ell)}}{s_i} \leq 1. \tag{A.24}$$

We now examine this threshold in greater detail. Recall from Equation (A.15) that

$$s_i = d_i + (1-d_i)c_i$$

where $c_i$ constitutes the corrected nIBD-to-IBS model. We can therefore interpret $s_i$ as the expected IBS sharing at locus $i$, predicated on the average relatedness $d_i$ at locus $i$ in the parasite sample. As such, the observable threshold

$$\frac{1}{m}\sum_{i=1}^{m}\frac{S_i^{(k,\ell)}}{s_i} = 1 \tag{A.25}$$

can be thought to stratify pairs with below-average vs above-average relatedness. We interpret the LHS of Equation (A.25) as a weighted measure of pairwise IBS sharing, with pairwise IBS at loci with a limited sample proportion of IBS pairs (i.e., small $s_i$) weighted more heavily.

Our interpretation of the MLE $\hat{r}^{(k,\ell)}$, mirroring Weir and Goudet [21], is thus two-fold:

- $\hat{r}^{(k,\ell)} = 0$ indicates that the parasite pair $(k, \ell)$ has below-average relatedness relative to the parasite sample, but is otherwise uninformative (i.e., does not quantify the extent to which pairwise relatedness deviates from the average relatedness in the parasite sample).
- $\hat{r}^{(k,\ell)} > 0$ quantifies the extent to which relatedness between the parasite pair $(k, \ell)$ exceeds the average relatedness in the parasite sample, or the amount of relatedness between the parasite pair $(k, \ell)$ that cannot be explained by average relatedness alone.

The adjustment for 'average relatedness' in the parasite sample is predicated on the locuswise sample proportion of IBD pairs $d_i$. If $d_i$ is taken to be constant across all loci $i = 1, \ldots, m$, then from Equations (A.20) and (A.22), a *non-zero* MLE $\hat{r}^{(k,\ell)}$ can be written in the form

$$\hat{r}^{(k,\ell)} = \frac{\hat{r}^{(k,\ell)}_{\text{corrected}} - d}{1 - d}$$

where $\hat{r}^{(k,\ell)}_{\text{corrected}}$ is the MLE obtained under the corrected nIBD-to-IBS model $c_i$ and the index $i$ is dropped on account of all $d_i$ being equal. A schematic of the relationship between $\hat{r}^{(k,\ell)}_{\text{corrected}}$ and $\hat{r}^{(k,\ell)}$ is shown in Figure A.4. Accounting for the variability of $d_i$ across loci, we predict, would yield a fuzzy elbow-like characteristic, with a change point near the average sample relatedness

$$\bar{d} = \frac{1}{m}\sum_{i=1}^{m} d_i.$$

For a sample of individuals with a positively-skewed distribution of pairwise relatedness, we would expect the majority of parasite pairs to exhibit "below-average" relatedness, manifest in pronounced zero inflation of the pairwise MLEs $\hat{r}^{(k,\ell)}$.

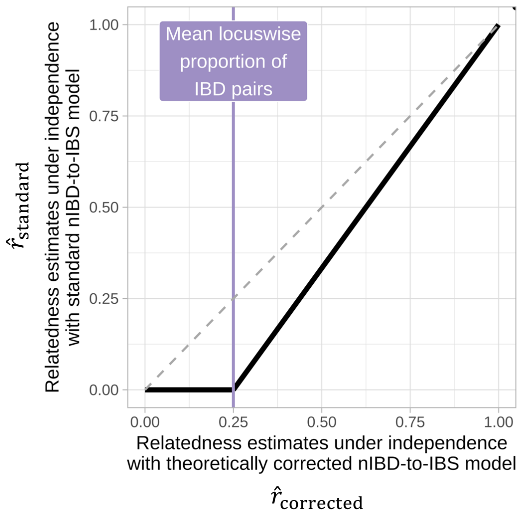

**Figure A.4:** Schematic of MLEs of pairwise relatedness under the model of (n)IBD independence coupled with the standard (misspecified) model $\hat{r}_{\text{standard}}$ vs corrected nIBD-to-IBS model $\hat{r}_{\text{corrected}}$.

#### A.3 Numerical results

The theoretical results derived in Sections A.2.1.2, A.2.2.1, and A.2.2.2 all support the central idea that relatedness is systematically underestimated when the independence model of relatedness is coupled with standard (n)IBD-to-observational models, due to the partial encoding of average population relatedness in the sums of squares of sample allele frequencies. Here, we present various numerical results that complement our theoretical analysis (Section A.3.1). We also interrogate the consequences of marker density and linkage, motivated by empirical data pointing towards systematic differences between pairwise relatedness estimates generated using sparse and WGS data: most strikingly, the pronounced zero inflation of sparse-data estimates relative to WGS estimates (see Figure S2Q of Taylor et al. [20] for a comparison of relatedness estimates generated using a 93-SNP molecular barcode vs 34911 WGS SNPs for a parasite sample from the Thai-Myanmar border). As an explanation, we suggest that exploiting linkage structure within dense data using the HMM of relatedness mitigates underestimation driven by standard observation models (Section A.3.3). Zero-inflation in relatedness estimates, arising from an over-reliance on the underlying (n)IBD-to-observational model, can be explained through comparative plots of IBS sharing (Section A.3.4). In light of these results, we propose a practical diagnostic to gauge the average locuswise relatedness $\bar{d}$ in a sample and approximate the severity of underestimation, predicated on the comparison of relatedness estimates under the independence model vs HMM for dense data (Section A.3.5). Zero inflation renders mean($\hat{r}$) under (n)IBD independence a poor approximation of $\bar{d}$.

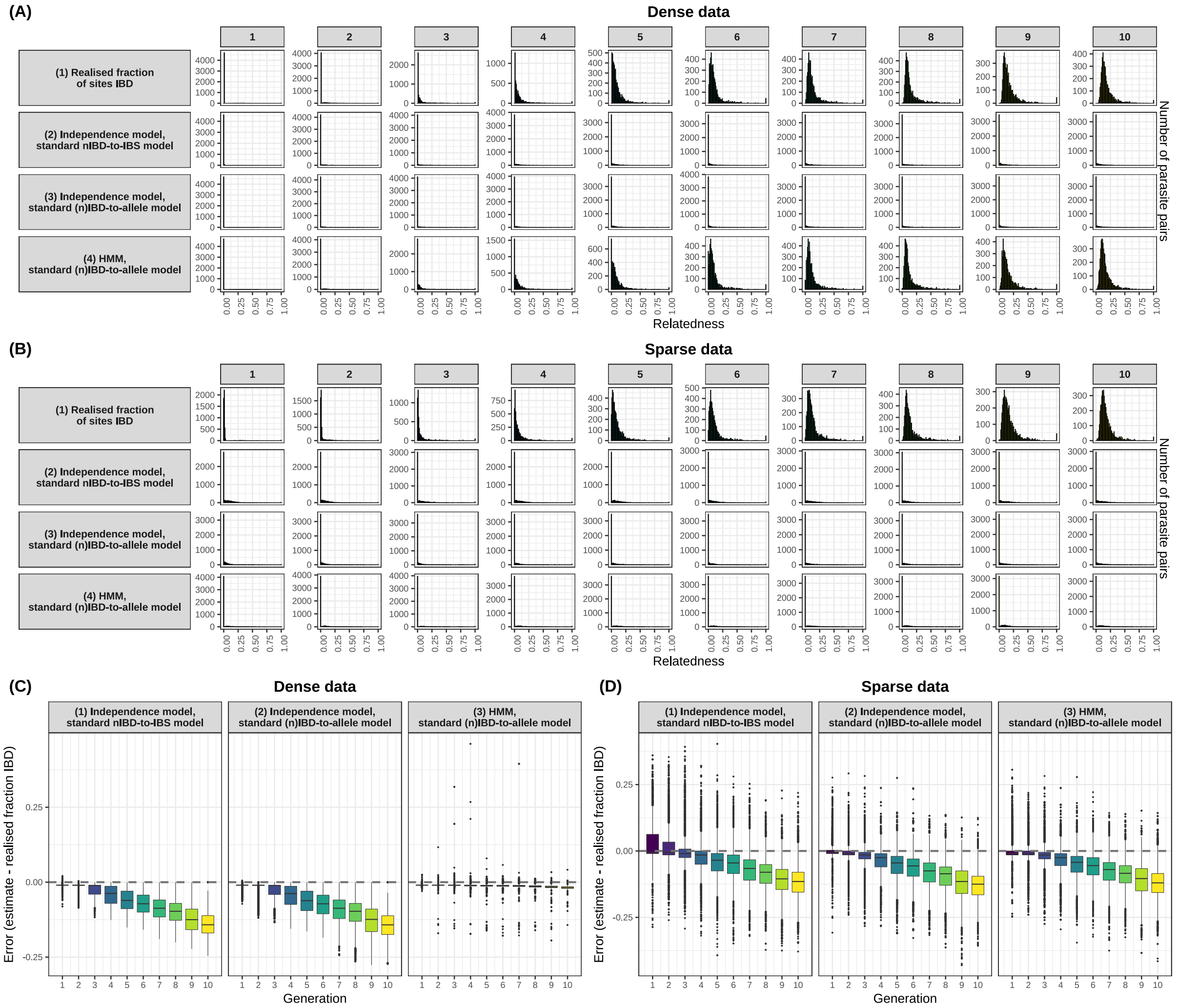

**Figure A.5:** Pairwise relatedness estimates predicated on the standard sample-allele frequency based models and either allelic states or IBS descriptives; either the independence model of relatedness vs HMM; and either dense or sparse data. Error, where applicable, is measured relative to the realised fraction of polymorphic sites IBD for each pairwise comparison. Sparse data have been obtained by selecting 200 polymorphic markers uniformly at random without replacement from the dense simulated dataset. Results are based a single realisation of the simulation model, with parameter values as per Table B.1.

##### A.3.1 Relatedness is systematically underestimated using standard models

Relatedness is systematically underestimated using standard models (Figure A.5). This holds whether observations are IBS descriptives (plots C1, D1) or alleles (plots C2–C3, D2–D3), whether the standard model assumes (n)IBD independence (plots C1–C2, D1–D2) or not (plots C3, D3), and whether markers are dense or sparse (plots C and D, respectively). Estimates are severely zero-inflated when data are sparse (plots B2–B4) and when dense data are fit under (n)IBD independence (plots A2–A3). Less severe zero-inflation and underestimation when the HMM is fit to dense data (plots A4 and C3, respectively) is addressed in Sections A.3.3 and A.3.4.

##### A.3.2 Underestimation using standard models is due to the partial encoding of population relatedness in sample allele frequencies

The degree of underestimation of pairwise relatedness using standard models increases with the number of generations of inbreeding because the standard nIBD-to-nIBS model is increasingly misspecified, in that the sums of squares of sample allele frequencies encode more relatedness (Figure A.6). The unbiased nature of relatedness estimates generated under the corrected independence model fit to (n)IBS observations (Figure A.7) also supports, by contrast, the idea that underestimation is due to the use of sample allele frequencies within standard observation models. This is true of both dense and sparse data, although estimates based on sparse data are less precise.

##### A.3.3 Relatedness is less severely underestimated using the standard HMM fit to dense data

Consider an inbred population with elevated relatedness (Figure A.8A). Increasing the marker count while accounting for linkage (estimation under the HMM with the standard (n)IBD-to-allele model) generates more precise and less biased estimates (Figure A.8B). Meanwhile, increasing the marker count without accounting for linkage (estimation under (n)IBD independence with either the standard nIBD-to-IBS or (n)IBD-to-allele model) does not remove bias (Figure A.8C, D). We believe this is because estimation under the HMM exploits linkage information, which increases with marker density. Meanwhile, estimation under the independence model does not exploit linkage information, regardless of its extent in the data. Instead, the independence model is entirely reliant on sample allele frequencies, which render the observational model misspecified. Otherwise stated, although sample allele frequencies are used under

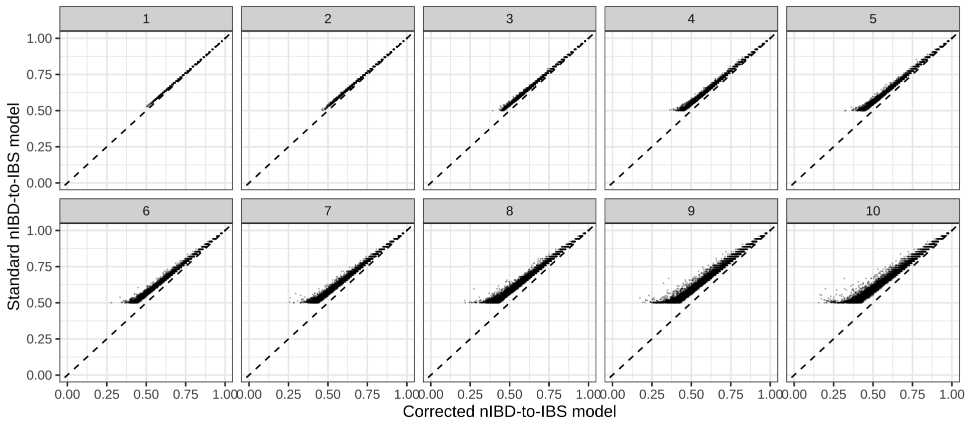

**Figure A.6:** The sample proportion of nIBD pairs that are IBS $c_i$ (i.e., the corrected nIBD-to-IBS model, accounting for the locuswise average relatedness) vs the locuswise proportion of IBS pairs $s_i$ (i.e., the standard nIBD-to-IBS model, based on sample allele frequencies) across successive generations of inbreeding for a single realisation of the simulation model, with parameter values as per Table B.1. For biallelic markers, it is necessarily the case that $s_i \geq 0.5$; however, $c_i$ may lie anywhere in the range $[0, 1]$.

both the standard HMM and independence model, the standard HMM is less reliant upon them and thus less susceptible to the misspecification they cause.

***Aside***: Under the HMM, the extent to which pairwise relatedness is underestimated decreases over the zero to one range granted data are sufficiently dense to encode linkage structure (Figure A.9). This is because, for a given marker count, the extent of linkage depends on the length of shared IBD segments, which is greater for recent relatives, and thus correlated with relatedness. We mention this as an aside only, because linkage due to recent ancestry is an inherent property of a given parasite genotype pair, not something we can control.

##### A.3.4 Comparative plots of IBS distributions capture the extent of zero-inflation under the independence model

The persistence of zero inflation in dense-data relatedness estimates under the independence model can be explained through comparative plots of IBS distributions.

For a given pair of individuals, the fraction of (polymorphic) sites that are IBS can be computed. The computation can then be repeated for all possible pairs. Hereafter, we refer to this set of values as the empirical IBS distribution for all pairs. As a comparator, under the standard nIBD-to-IBS model, i.e., $\boldsymbol{s} = (s_1, \ldots, s_{n_{\text{markers}}})$, the expected IBS distribution for ostensibly unrelated parasite pairs ($r = 0$) comprises a rescaled Poisson binomial distribution (i.e., a sum

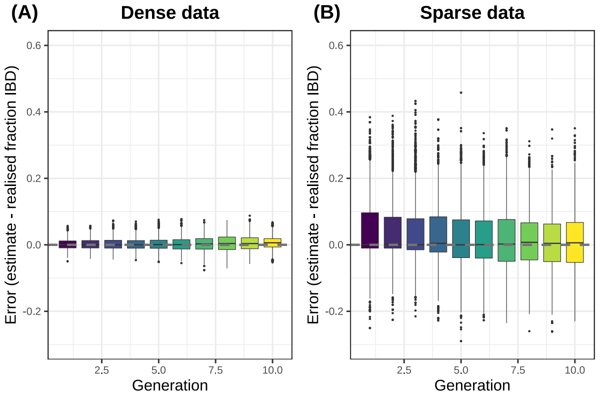

**Figure A.7:** Relatedness estimates predicated on the independence model and the corrected locuswise nIBD-to-IBS model $c_i$. Sparse data have been obtained by selecting 200 polymorphic markers uniformly at random without replacement from the dense simulated dataset. Results are based a single realisation of the simulation model, with parameter values as per Table B.1.

of independent, but not necessarily identically-distributed Bernoulli random variables):

$$Y_0(\boldsymbol{s}) = \frac{1}{n_{\text{markers}}} \sum_{i=1}^{n_{\text{markers}}} X_i(s_i)$$

where

$$X_i(s_i) \overset{\text{independent}}{\sim} \text{Bernoulli}\big(s_i\big).$$

$Y_0(\boldsymbol{s})$ is also applicable under the HMM, because the HMM collapses down to the independence model for unrelated pairs.

Comparison of the empirical IBS distribution for all pairs and the expected IBS distribution for ostensibly unrelated pairs (Figure A.10A) shows that there is a range of empirical IBS values less that the expected IBS distribution for ostensibly unrelated pairs. The majority of these pairs have zero-valued relatedness estimates under the independence model, because the independence model is entirely reliant on the standard nIBD-to-IBS model (Figure A.10B, orange). Otherwise stated, the extent to which empirical IBS values fall below the expected IBS distribution for unrelated pairs corresponds to the extent of zero-inflation under independence. This result is consistent with the notion that locuswise average relatedness is partially encoded within the locuswise sample proportion of IBS pairs $s_i$, whereby the expected IBS distribution for ostensibly unrelated pairs ought to be reinterpreted as the expected IBS distribution for pairs with 'average' relatedness. Consequently, a range of below-average relatedness values map onto zero-valued relatedness estimates under the independence model. Meanwhile, they have small but non-zero estimates under the HMM model, because the HMM is less reliant on the

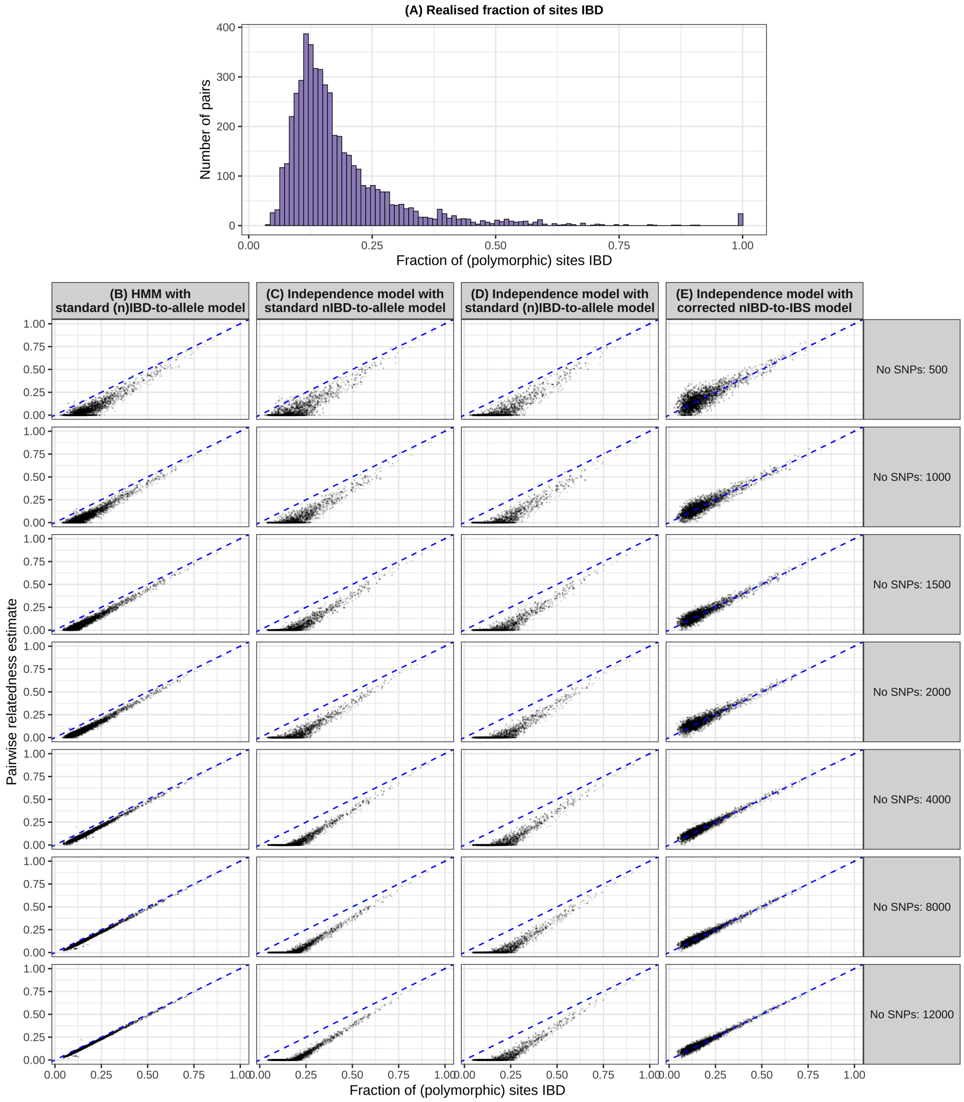

**Figure A.8:** Summary of pairwise relatedness for simulated data after 10 generations of inbreeding, yielding the distribution of realised relatedness (i.e., the fraction of simulated polymorphic sites that are IBD for each parasite pair) shown in (A). For downsampled marker subsets (selected uniformly at random without replacement over the set of polymorphic markers), realised relatedness is calculated over the marker subset and treated as the ground truth. We compare realised relatedness against pairwise relatedness estimates generated under (B): the HMM with the standard (n)IBD-to-allele model; (C): (n)IBD independence with the standard (n)IBD-to-allele model; (D): (n)IBD independence with the standard nIBD-to-IBS model; (E): (n)IBD independence with the theoretically-corrected nIBD-to-IBS model. Results are based on a single realisation of the simulation model, with parameter values as per Table B.1.

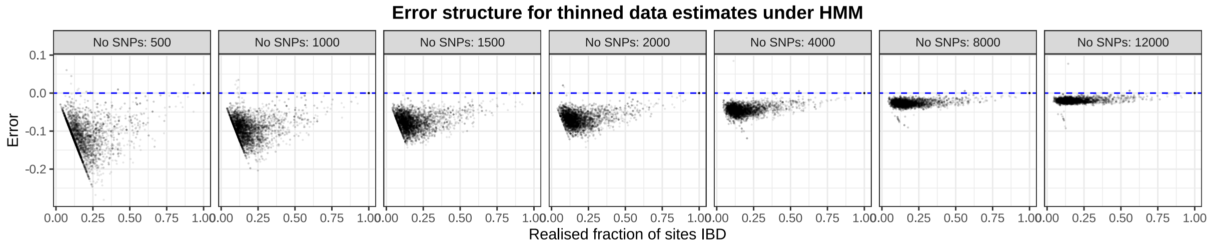

**Figure A.9:** Error structure for relatedness estimates of downsampled marker subsets (selected uniformly at random without replacement over the set of polymorphic markers) at generation 10 under either the HMM of relatedness with the standard (n)IBD-to-allele model, compared to the realised fraction of polymorphic sites IBD for each parasite pair. Results are based on a single realisation of the simulation model, with parameter values as per Table B.1.

underlying (n)IBD-to-observation model (Figure A.10B, green). This implies that strategies that focus on high minor allele frequencies will have less power to resolve population structure (see [25]).

##### A.3.5 Dense-dense data elbow-like plots approximately capture the extent of underestimation

We leverage the precision of estimates generated using dense data to design a practical diagnostic for identifying an approximate of the corrective value $\bar{d}$. More specifically, although relatedness estimates generated under the HMM using dense marker data from inbred populations are underestimates (Figures A.8 and A.9), they are sufficiently less biased downwards compared with those generated under independence to reveal an elbow-like pattern (Figure A.11B), which is otherwise only accessible using either simulation (Figure A.11A) or theory (Figure A.4). The use of dense marker data under the independence helps to resolve the pattern, because estimates based on dense data are more precise.

#### A.4 Empirical results

We show, using real data, that the theoretical and numerical results apply in real life. To do so, we perform a case study of an inbred parasite sample from Guyana, in which population structure is easy to diagnose, and for which WGS SNP data are available. For a parasite sample derived from a single population, we chart out a practical roadmap for assessing systematic bias and the effects of marker sparsity in the main text. In this context, the WGS SNP data serve two purposes: to gauge the average population-relatedness, and consequently the severity of

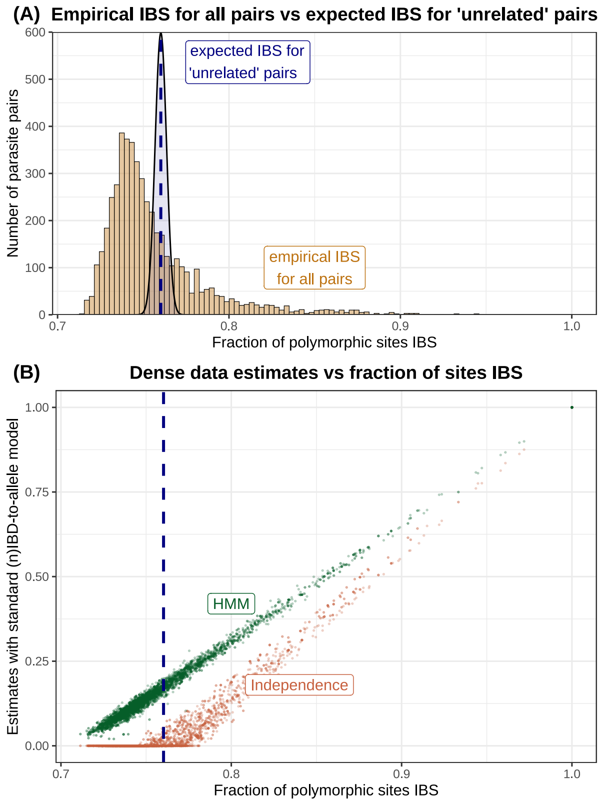

**Figure A.10:** For dense data at generation 10:

(A) The empirical pairwise distribution of IBS sharing for all pairs (ochre), defined to be the fraction of polymorphic sites that are IBS, compared to the expected IBS distribution $Y_0$ for ostensibly unrelated pairs (blue).

(B) Relatedness estimates generated under the HMM (green) or independence model (orange) using the standard (n)IBD-to-allele model vs the fraction of polymorphic sites IBS.

Results are based on a single realisation of the simulation model, with parameter values as per Table B.1. The probability mass function for the Poisson binomial distribution has been computed using the R package `poisbinom` [19].

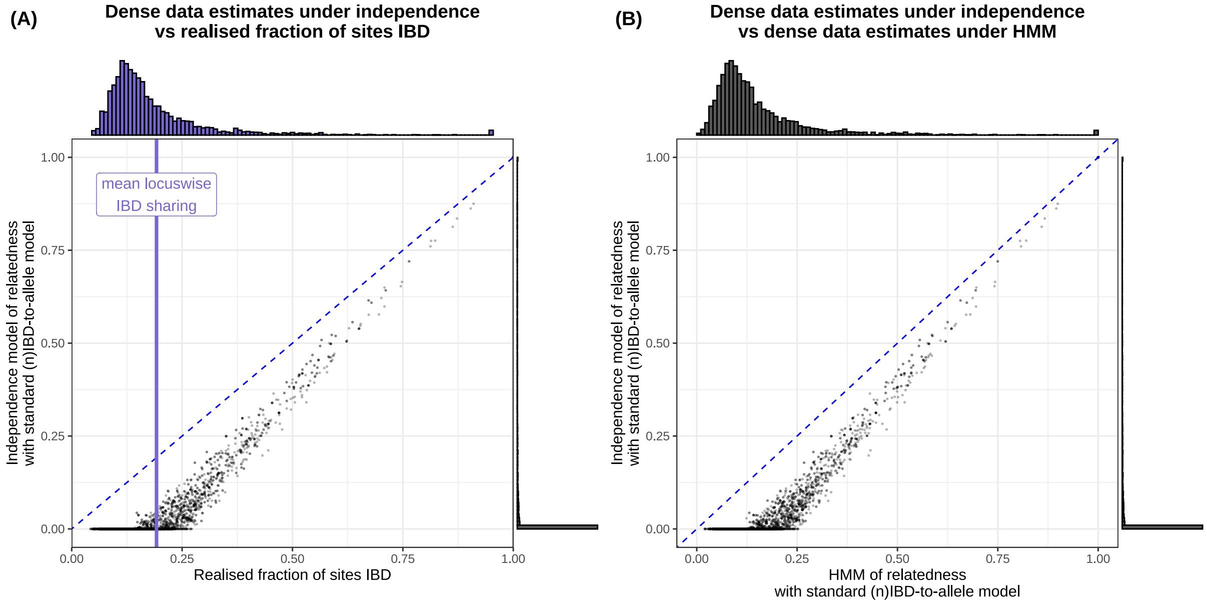

**Figure A.11:** Relatedness estimates for dense genotypic data at generation 10 under the independence model vs (A) the realised fraction of polymorphic sites IBD or (B) corresponding estimates under the HMM of relatedness. Here, we use the standard (n)IBD-to-allele model. Results are based on a single realisation of the simulation model, with parameter values as per Table B.1.

underestimation; and to gauge the marker densities required to answer questions of practical interest, particularly for threshold-based IBD analyses.

We devote the present Appendix A.4 to the effect of population structure, unconsidered hitherto, under which our theoretical and numerical results do not necessarily hold. To do so, we pool the parasite sample from Guyana ($n = 278$ isolates) [36] with an additional $n = 28$ high-quality isolates from Colombia [33], yielding genotypes at $n = 30694$ (polymorphic) biallelic SNPs. Population structure appears to be pronounced: principal coordinates analysis (PCoA), based on pairwise fractions of IBS markers, yields two distinct clusters, stratified by country (Figure A.12).

##### A.4.1 Diagnostics of population structure

We propose two diagnostics to screen for population structure. First we examine the empirical distribution of pairwise fractions of IBS markers. In the absence of population structure, we would expect to see a unimodal IBS distribution. The multimodal IBS distribution shown in Figure A.13, in contrast, is indicative of population structure [30]: we view it as a mixture distribution, with components stratified by within- and between-subpopulation comparisons.

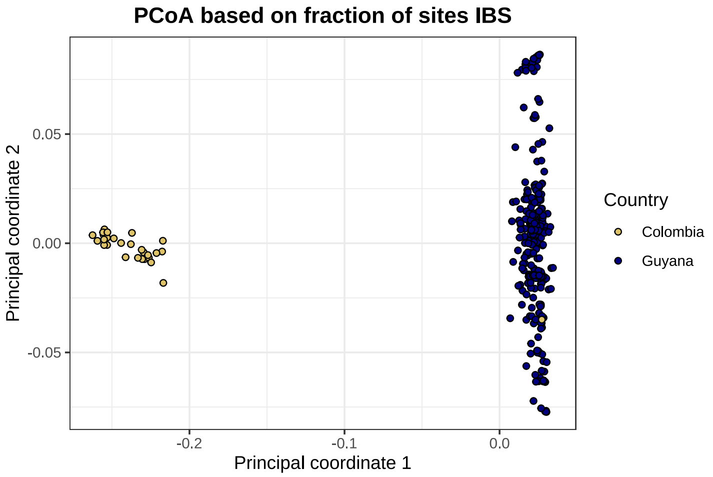

**Figure A.12:** Principal coordinates analysis, with a distance matrix based on pairwise fractions of IBS markers. The major subpopulation (navy blue) is defined by a value of at least -0.1 for principal coordinate 1. PCoA has been performed using the R function `stats::cmdscale` [31].

The well-separated components suggest pronounced differentiation between allele frequencies in the constituent subpopulations.

An alternative diagnostic, linking back to our characterisation of observation models predicated on allelic states vs IBS descriptives in Section A.2.1, is the comparison of relatedness estimates under (n)IBD independence predicated on the standard (n)IBD-to-allele model vs the standard nIBD-to-IBS model (Figure A.14). Allelic states are more informative than IBS descriptives. Under the (n)IBD-to-allele model, a shared minor allele points towards IBD more strongly than a shared major allele; this relative weighting is erased when we shift to IBS descriptives. In the absence of population structure, we would expect each pair of individuals to share a mixture of

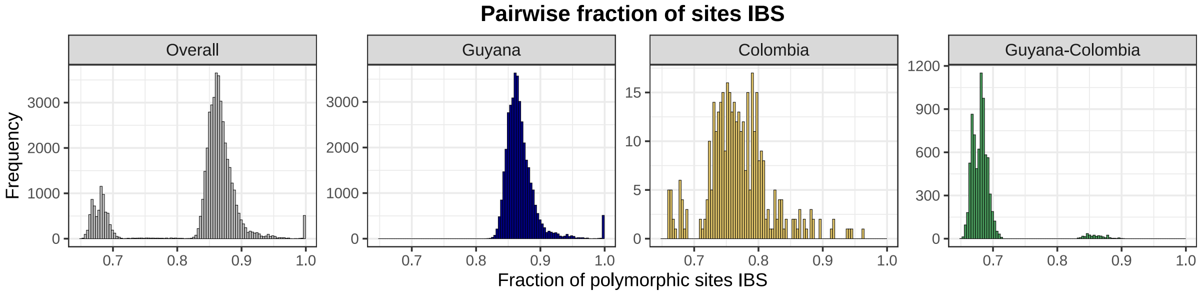

**Figure A.13:** Pairwise fractions of IBS markers as a diagnostic of population structure. In light of mising data, IBS sharing for each pair is defined to be the fraction of polymorphic, comparable (i.e., non-missing) sites that are IBS. We stratify comparisons within and between the subpopulations defined in Figure A.12 using PCoA.

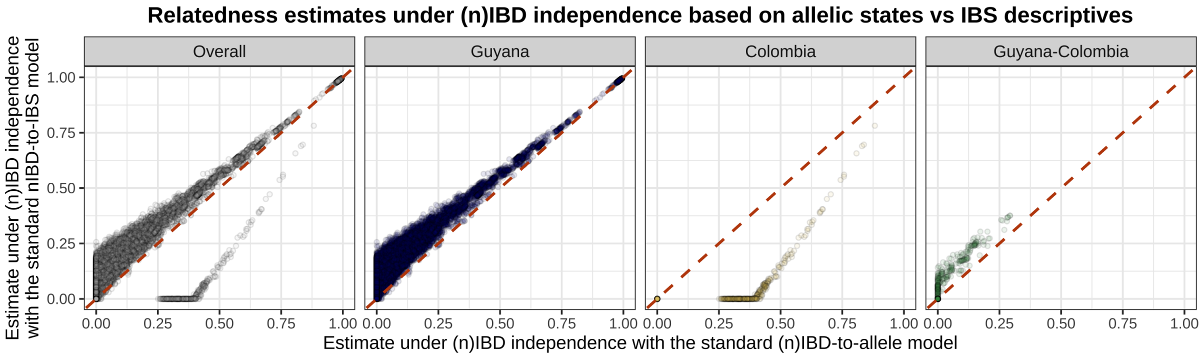

**Figure A.14:** Comparison of relatedness estimates generated under (n)IBD independence using the standard nIBD-to-IBS model vs the standard (n)IBD-to-allele model as a diagnostic of population structure. We stratify comparisons within and between the subpopulations defined in Figure A.12 using PCoA.

major and minor alleles; a shift from allelic states to IBS descriptives would therefore introduce noise, but we would expect the over/under-weighting of shared major/minor alleles to average out across markers. We would, however, expect systematic differences to emerge in the presence of population structure. Comparisons within the major subpopulation, for instance, systematically yield parasite pairs with a shared major allele, yielding higher relatedness estimates based on IBS descriptives relative to allelic states; the converse applies to the minor subpopulation. Systematic patterns akin to those in Figure A.14 are thus indicative of population structure.

##### A.4.2 Dense data diagnostic in the presence of population structure

We now examine the dense data diagnostic proposed in Section A.3.5 — namely, a comparative plot of relatedness estimates under the independence model vs HMM for a dense WGS SNP dataset — in the presence of population structure. In the absence of population structure (main text), we observe a clean elbow-like pattern concordant with our theoretical predictions (Figure A.4) and numerical results (Figure A.11). Population structure, however, yields the emergence of multiple elbows, corresponding to different within- and across-population comparisons, with a different structure depending on whether estimates under (n)IBD independence are predicated on allelic states or IBS descriptives (Figure A.15). Since sample allele frequencies represent a weighted average across subpopulations, interpretation of the branch points is unclear. The cross-population observation model proposed by Schaffner et al. [23], in which sample allele frequencies are stratified by subpopulation, may yield more interpretable results.

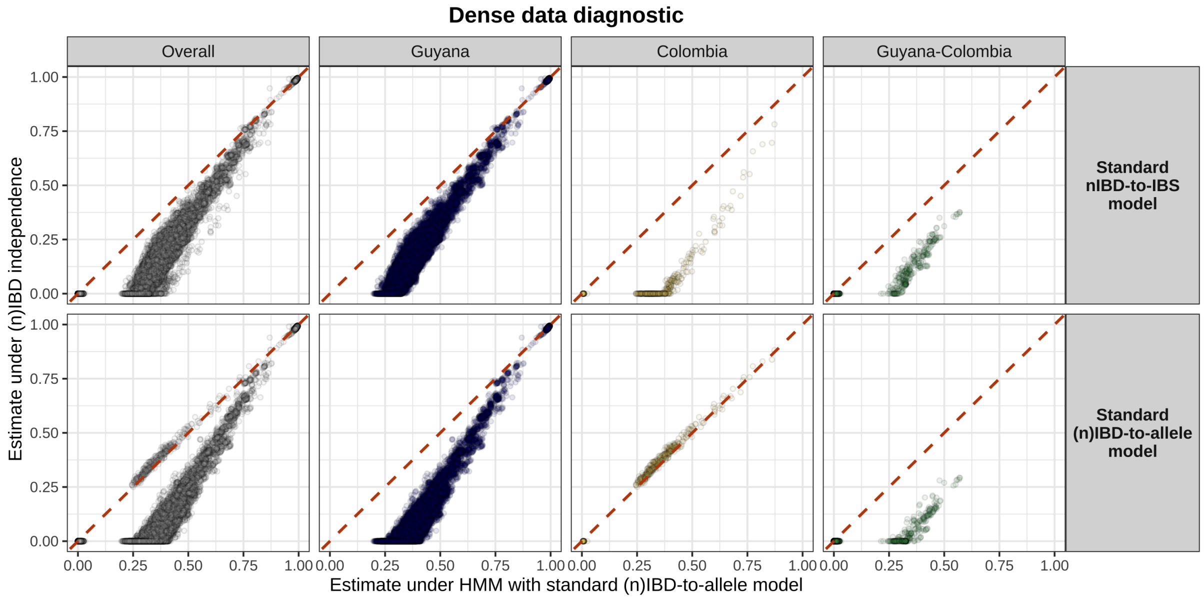

**Figure A.15:** Dense data diagnostic, comparing relatedness estimates under the HMM and standard (n)IBD-to-allele model vs (n)IBD independence. We stratify comparisons within and between the sub-populations defined in Figure A.12 using PCoA.

##### A.4.3 Zero inflation and population structure

The nIBD-to-IBS model leads to the systematic over/under-weighting of shared major/minor alleles relative to the (n)IBD-to-allele model. When sample allele frequencies are averaged over several subpopulations and estimates are generated under (n)IBD independence, allelic states yield zero inflation in comparisons within the major subpopulation; while IBS descriptives yield zero inflation in comparisons within the minor subpopulation (Figure A.16). Theoretically-predicted and numerically-validated results pertaining to zero inflation, therefore, are sensitive to population structure.

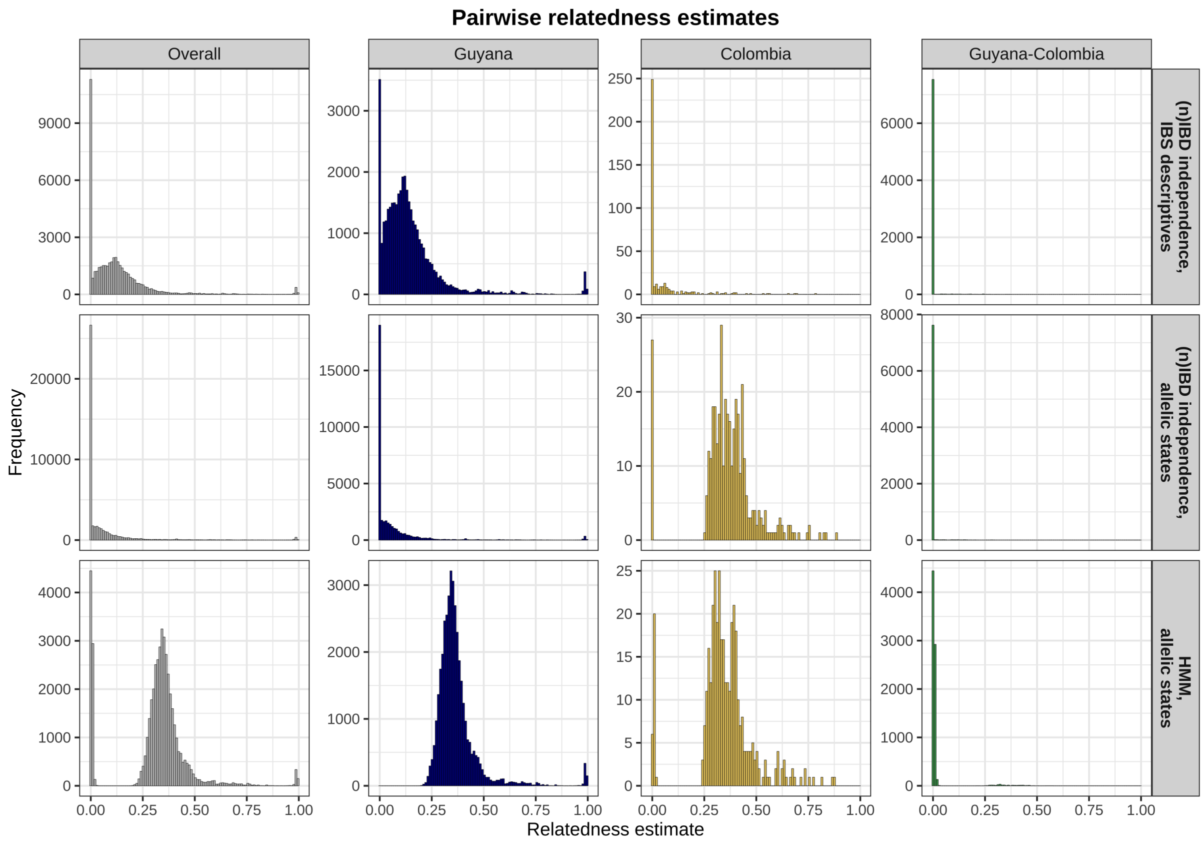

**Figure A.16:** Pairwise relatedness estimated under (n)IBD independence with the standard nIBD-to-IBS model; (n)IBD independence with the standard (n)IBD-to-allele model; and the HMM with the standard (n)IBD-to-allele model. We stratify comparisons within and between the subpopulations defined in Figure A.12 using PCoA.

### Appendix B

### Simulation model

We describe a simulation model that starts with a population whose ancestry is defined in terms of a bygone outbred founder population and then simulates successive generations of inbreeding. Our model differs from the canonical Wright-Fisher model of stochastic drift, which concerns a single locus [15], on three grounds:

- We simulate multiple loci and introduce linkage between them through a discrete-time homogeneous Markov chain.
- We introduce clonal and sibling substructure through relationship graphs.
- On top of the sibling and clonal substructure, we introduce an increased propensity for selfing and sibling-sibling mating through a single parameter; in doing so, we capture departure from panmixia. In reality, a single parameter cannot represent multiple processes that govern selfing and sibling-sibling mating between malaria parasites: selfing between malaria parasites is always viable theoretically; given a monoclonal mosquito infection, it is inevitable. Mosquito-to-human-to-mosquito cotransmission enhances the probability of sibling-sibling mating [32].

Demographic processes like immigration, mutation and selection are not accounted for.

Our description is structured as follows. In Section B.1, we introduce a model under which offspring are generated as a mosaic of two parents, allowing for linkage between markers which are all equidistant. We then propose a model for our "generation zero" population, whose ancestry is formulated in terms of a bygone outbred founder population (Section B.2). In Section B.3, we propose a model under which the population in the present generation guides the generation of the population in the next generation. Designed to capture a single generation of inbreeding, our framework is underpinned by the relationship graph model of Taylor et al. [27]. In Section B.4, we detail our approach for assessing misspecification under the standard nIBD-to-IBS model. A summary of simulation parameters, and their respective tradeoffs, is provided in Section B.5.

#### B.1 Generating a single individual with inter-marker linkage

We consider some number, $n_{\text{markers}}$, of multiallelic, *equidistant* markers, spanning a single chromosome, and indexed $i = 1, \ldots, n_{\text{markers}}$, with each marker treated as a point polymorphism [26]. Each time we simulate a single meiosis, we generate a single offspring as a random $n_{\text{markers}}$-mosaic of parents that are labelled $\{a_1, a_2\}$ respectively. In reality, a single meiosis between two parental genotypes generates four meiotic offspring, which then replicate asexually into thousands of sporozoites; on average, meiotic offspring are related to each other by 1/3 and to each parent by 1/2 if parental genotypes are unrelated [24, 27].

Denote by

$$\boldsymbol{b} = \big(b_1, \ldots, b_{n_{\text{markers}}}\big)$$

the mosaic of an offspring of parents $\{a_1, a_2\}$, with $b_i = a_1$ if marker $i$ is inherited from parent $a_1$ and $b_i = a_2$ otherwise.

Assuming a genomewide constant recombination rate of $\rho$, $\boldsymbol{b}$ is governed by a discrete-time, homogeneous Markov chain with transition matrix

$$\begin{pmatrix} \mathbb{P}(b_{i+1} = a_1 | b_i = a_1) & \mathbb{P}(b_{i+1} = a_1 | b_i = a_2) \\ \mathbb{P}(b_{i+1} = a_2 | b_i = a_1) & \mathbb{P}(b_{i+1} = a_2 | b_i = a_2) \end{pmatrix} = \begin{pmatrix} 0.5\big(1 + e^{-\rho d}\big) & 0.5\big(1 - e^{-\rho d}\big) \\ 0.5\big(1 - 0.e^{-\rho d}\big) & 0.5\big(1 + e^{-\rho d}\big) \end{pmatrix},$$

equivalent to the Markov model of relatedness (Equation (A.3)) in the case $r = 0.5$ with switch rate $\kappa = 1$ where $d$ is measured in base pairs (bp) and the recombination rate $\rho$ has units M/bp.

Due to discrete sampling of the genome, consecutive markers inherited from the same parent may be separated by an even number of recombination breakpoints, which occur between base pairs. For the purposes of simulating ancestry at a fixed set of markers, we do not distinguish whether or not a string of markers inherited from the same parent is separated by recombination breakpoints (Figure B.1), and assume that a Poisson process governs the position of recombination breakpoints [4].

Observe that

$$\mathbb{P}\left(b_{i+1} = b_{i+2} = \ldots b_{i+m-1} = a_1, b_{i+m} = a_2 \,\middle|\, b_i = a_1\right) = \frac{1}{2^m}(1 - e^{-\rho d})\big[1 + e^{-\rho d})\big]^{m-1}.$$

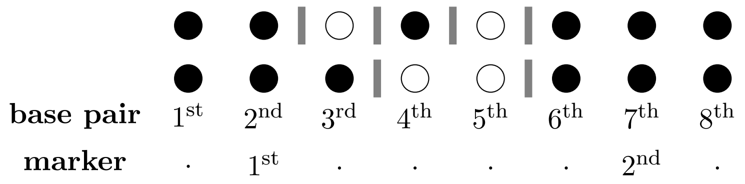

**Figure B.1:** Two examples of pairwise (n)IBD states across a stretch of 8bp. Recombination breakpoints (vertical bars) are shown between IBD (black) and nIBD (white) loci. Consecutive IBD markers may be separated by an even number of recombination breakpoints, that may not be detectable due to the discrete sampling of the genome.

If we ignore (temporarily) the finite number of markers $n_{\text{markers}}$, then given $b_i = a_1$, the number of consecutive markers (from $i$ inclusive) inherited from parent $a_1$ is geometrically distributed with state space $\mathbb{N}$ and mean length

$$\bar{M} := \frac{2}{1 - e^{-\rho d}}.$$

Stretches of consecutive markers inherited from parent $a_2$ are identically-distributed.

We can therefore treat $\boldsymbol{b}$ as a truncated alternating renewal process, where segments alternate from each parent and are independent and identically distributed (i.i.d.) with length

$$L \sim \text{Geometric}\Big(\frac{1}{\bar{M}}\Big),$$

where the geometric distribution is taken to have support $\mathbb{N}$. The case $\bar{M} = 2$ reduces to the independence model.

To simulate an offspring under this model, we sample a parent $a_1$ or $a_2$ with probability 1/2; sample a geometric segment length; select the alternative parent and re-sample a geometric segment length; string together alternating geometric segments from each parent; and then truncate once a length of $n_{\text{markers}}$ has been obtained, to account for the finite length that is ignored above. An alternative parametrisation with i.i.d. geometric segments of mean length $\bar{B} = \bar{M}/2$ would yield a model of the number of consecutive markers inherited from a single parent before a recombination breakpoint. At every junction, we would then need to select a new parent with probability 1/2, rather than switching deterministically between parents.

#### B.2 Generating a population of individuals at generation zero

We now propose a model under which to generate a "generation zero" population whose ancestry is constructed in terms of a bygone outbred founder population. This construction allows us to distribute background relatedness across the generation zero population.

Firstly, let us consider a founder population comprising some number of unrelated equifrequent founders, $n_{\text{founders}}$. At each marker $i = 1, \ldots, n_{\text{markers}}$, we select the allele $q \in \{1, \ldots, y\}$ with probability $p_i(q)$

$$A_{\text{founder}}(f, i) \overset{i.i.d.}{\sim} \text{Categorical}\big(p_i(1), \ldots, p_i(y)\big)$$

independently for each founder $f = 1, \ldots, n_{\text{founders}}$. Here, multiallelic markers are treated as point polymorphisms [26]. For large $n_{\text{markers}}$, each founder is likely to harbour a unique allelic sequence.

We represent individuals within subsequent generations as mosaics of the $n_{\text{founders}}$ founders; that is the ancestry of individual $j$ in generation $k$ is represented in the form

$$\boldsymbol{F}(j,k) = \big(f_1, \ldots, f_{n_{\text{markers}}}\big) \in \big\{1, \ldots, n_{\text{founders}}\big\}^{n_{\text{markers}}}.$$

Barring genotyping error, the vector of allelic states observed for individual $j$ in generation $k$ is then given by

$$\boldsymbol{A}(j,k) = \big(A_{\text{founder}}(f_1, 1) \ldots, A_{\text{founder}}(f_{n_{\text{markers}}}, n_{\text{markers}})\big).$$

The ancestry of each individual $j$ in generation zero is independently sampled uniformly at random from the set of all founder sequences, that is,

$$\boldsymbol{F}(j,0) \overset{\text{i.i.d.}}{\sim} \text{Uniform}\big[\{1,, \ldots, n_{\text{founders}}\}^{n_{\text{markers}}}\big], \tag{B.1}$$

and we initialise a population relationship graph for generation zero by placing stranger edges between all individuals/nodes. Under this model, low-level background relatedness is maximally spread across parasite pairs in generation zero. Implicit in this construction is the assumption that there is sufficient temporal separation between the founder population and generation zero to break down inter-marker linkage. If we instead modelled generation zero as one generation of mating from the founder population, relatedness would be concentrated on a small subset of pairs with shared parents, consistent with recent rather than background relatedness (Section B.5.1), and Equation (B.1) would not hold because each individual in generation zero would be constructed as a mosaic of at most two founders.

***Aside***: The uniform assumption (Equation (B.1)) is also compatible with an (n)IBD-to-allele model predicated on founder allele frequencies. Because Equation (B.1) holds, the probability of randomly sampling the allelic sequence

$$\mathbf{A} = (q_1, \ldots, q_{n_{\text{markers}}})$$

for each individual in generation zero is proportional to the number of ways of recovering that sequence as a mosaic of founders:

$$\mathbb{P}(\boldsymbol{A}) = \frac{1}{(n_{\text{founders}})^{n_{\text{markers}}}} \sum_{f_1=1}^{n_{\text{founders}}} \cdots \sum_{f_{n_{\text{markers}}}=1}^{n_{\text{founders}}} \prod_{i=1}^{n_{\text{markers}}} \mathbb{1}\{A_{\text{founder}}(f_i, i) = q_i\}. \tag{B.2}$$

Interchanging the summation and product in Equation (B.2), we can equivalently write

$$\mathbb{P}(\boldsymbol{A}) = \frac{1}{(n_{\text{founders}})^{n_{\text{markers}}}} \prod_{i=1}^{n_{\text{markers}}} \sum_{f=1}^{n_{\text{founders}}} \mathbb{1}\{A_{\text{founder}}(f, i) = q_i\} = \prod_{i=1}^{n_{\text{markers}}} \theta_{\text{founder}}(q, i), \tag{B.3}$$

since the frequency of allele $q$ at locus $i$ in the founder population is

$$\theta_{\text{founder}}(q, i) := \frac{1}{n_{\text{founders}}} \sum_{f=1}^{n_{\text{founders}}} \mathbb{1}\{A_{\text{founder}}(f, i) = q\}. \tag{B.4}$$

Equation (B.3) is precisely the likelihood of observing the allelic sequence $\boldsymbol{A}$ that we would obtain by plugging founder allele frequencies (B.4) into the standard nIBD-to-allele model under locus independence.

#### B.3 Generating populations of individuals over successive generations of inbreeding

From generation $k = 1$ onwards, we simulate successive discrete, non-overlapping generations of inbreeding. We capture stochastic drift, with the imposition of additional sibling/clonal substructure in the line with the formulation of Taylor et al. [27]; but ignore the phenomena of immigration, mutation and selection.

We assume that the population size $n_{\text{individuals}}$ remains fixed across generations. As stated above, the population relationship graph for generation zero has stranger edges between all individuals. To simulate the ancestry of generation $k \geq 1$, given that of generation $(k-1)$, we perform the following steps:

1. Simulate a population-level relationship graph for generation $k$, characterised by clonal, sibling and stranger edges (Section B.3.1).

2. Conditional on the structure of the relationship graphs for generations $(k-1)$ and $k$, formulate each individual in generation $k$ as a mosaic of parents from generation $(k-1)$, allowing for an enriched probability of sibling-sibling crosses and selfing over generations (Section B.3.1.1).

3. Recover the ancestry structure (relative to founders $f = 1, \ldots, n_{\text{founders}}$) for each individual in generation $k$ using the encoding $\boldsymbol{F}(j, k-1)$.

The simulation structure is informed by the work of Taylor et al. [27], and draws on the R package `Pv3Rs` [35]. Below, we describe the first two steps in detail.

##### B.3.1 Generating relationship graphs

For each generation $k \geq 1$, we simulate a transitive relationship graph with sibling, clonal and stranger edges, following the framework of Taylor et al. [26]. Relationship graphs, which are designed to capture one generation of inbreeding, are sampled independently for each generation.

The function `sample_RG`, implemented in the R package `Pv3Rs` [35], simulates relationship graphs uniformly at random over the space of all allowable transitive graphs with sibling/clonal/stranger edges. Due to computational constraints, rather than directly simulating a relationship graph for a complete population of individuals, size $n_{\text{individuals}}$, we simulate subgraphs of size $m_{\text{subgraph}}$ uniformly at random using the function `sample_RG` of `Pv3Rs` [35]; and then amalgamate these subgraphs, with stranger edges for all between-subgraph comparisons, to recover a population-level relationship graph for $n_{\text{individuals}}$. As a consequence of this construction, population-level relationship graphs are sampled over a non-uniform distribution over the space of allowable transitive graphs, that is biased towards those containing small, relatively balanced sub-clusters of clones and siblings: while each subgraph is of fixed size $m_{\text{subgraph}}$, each clonal/sibling component is at most of size $m_{\text{subgraph}}$. When sub-clusters are balanced, the assumption of conditional independence between allelic and IBD states that underpins the allelic observation model is justified (Figure A.2, Section A.2.1.1).

The process of simulating individuals within subgraphs and then amalgamating subgraphs could be viewed as simulating individuals within infected hosts and then amalgamating infected hosts. Under this interpretation, $m_{\text{subgraph}}$ is a fixed number of parasite genotypes per host. Of those parasite genotypes, all could be clonal, as in a monoclonal infection; all could be strangers, as in a balanced multiclonal infection resulting from superinfection without cotransmission; all could be sibling as in a multiclonal infection resulting from balanced cotransmission without strangers; etc. The frequencies of these different scenarios are governed by the distribution over transitive graphs (Figure B.2A), which is uniform for convenience.

Alternatively, we could view simulated individuals within subgraphs as successfully transmitted sporozoites per mosquito bite, and $m_{\text{subgraph}}$ as the genotype count per bite. Under this construction, cotransmission could be modelled through subgraphs derived by sampling parents from a previous subgraph, as captured below through the parameter $p_{\text{cotransmission}}$. Superinfection, which implies independent bites, would be modelled through subgraphs derived by sampling parents from the population at large. However, under this construction, the partition of bites and genotypes across humans is abstracted.

###### B.3.1.1 Generating ancestry and thus genotypes conditional on relationship graphs for the present and previous generation

Given relationship graphs for generations $(k-1)$ and $k$, we use Algorithm 1 to simulate the ancestry structure of generation $k$ relative to generation $(k-1)$; that is, we formulate individuals in generation $k$ as crosses between parents from generation $(k-1)$. Each sibling/clonal component in generation $k$ is independently assigned two parents from generation $(k-1)$. Siblings are treated as *independent* crosses of the same parental pair, and therefore have expected relatedness 1/2 [24, 27]. Clones are treated as the same cross of a given parental pair, rather than a single individual sampled from generation $(k-1)$.

For each sibling/clonal component in the relationship graph for generation $k$, parental pairs are selected under two possible schemes:

- With probability $p_{\text{cotransmission}}$, we sample a constituent subgraph of size $m_{\text{subgraph}}$ (that is, one of the $n_{\text{individuals}}/m_{\text{subgraph}}$ subgraphs we generated uniformly at random over the set of allowable transitive relationship graphs of size $m_{\text{subgraph}}$) uniformly at random in generation $(k-1)$; and then select two parents uniformly at random *without* replacement from that constituent subgraph, yielding an elevated probability of sibling-sibling crosses and selfing. While parents are guaranteed to be sampled from the same subgraph, inbreeding is not guaranteed because a subgraph can contain strangers.
- With probability $(1-p_{\text{cotransmission}})$, we sample two parents from generation $(k-1)$ uniformly at random *with* replacement, whereby parentals may be derived from the same subgraph but are more likely from different subgraphs.

This sampling scheme is designed to capture stochastic drift, with the enriched probability of selfing and sibling-sibling crosses acting as a proxy for monoclonal mosquito infections and serial cotransmission [24].

#### B.4 Simulation model application

We treat the realised fraction of polymorphic markers that are IBD

$$R(j_1, j_2, k) := \frac{1}{n_{\text{markers}}} \sum_{i=1}^{n_{\text{markers}}} \mathbb{1}\{F_i(j_1, k) = F_i(j_2, k)\}$$

**Algorithm 1:** Simulate one generation of recombination (based on function `sample_lineages` of R package `Pv3Rs` [35])

**Input:** transitive relationship graph $G_{\text{current}}$ for current generation with sibling/clonal/stranger edges
equifrequent parents $\ell = 1, \ldots, n_{\text{parents}}$
stratification of parents $L_{\text{parent}}(g) \subset \{1, \ldots, n_{\text{parents}}\}$ by parental subgraph $g \in \{1, \ldots, n_{\text{individuals}}/m_{\text{subgraph}}\}$
probability of cotransmission $p_{\text{cotransmission}}$
number of $n_{\text{markers}}$ at which individuals are genotyped
size of parasite population $n_{\text{individuals}}$ in subsequent generation

**Result:** ancestry matrix $\boldsymbol{M} \in \{1, \ldots, n_{\text{parents}}\}^{n_{\text{individuals}} \times n_{\text{markers}}}$ for subsequent generation

Delete all stranger edges in $G_{\text{current}}$;
Decompose $G_{\text{current}}$ into fully-connected sibling/clonal subgraphs $S_1, \ldots, S_{n_{\text{subgraph}}}$;
**for** $j \in \{1, \ldots, n_{subgraph}\}$ **do**
    Delete all sibling edges in $S_j$;
    Decompose $S_j$ into clonal components $C_1^j, \ldots, C_{s_j}^j$;
    **if** $u \sim U(0, 1) < p_{cotransmission}$ **then**
        Sample parental subgraph $g \in \{1, \ldots, n_{\text{individuals}}/m_{\text{subgraph}}\}$ uniformly at random;
        Sample $\ell_1, \ell_2 \in L_{\text{parent}}(g)$ uniformly at random, without replacement;
    **else**
        Sample $\ell_1, \ell_2 \in \{1, \ldots, n_{\text{founders}}\}$ uniformly at random, with replacement;
    **for** $n \in \{1, \ldots, s_j\}$ **do**
        Simulate one meiosis $\boldsymbol{\ell}_{\text{offspring}} \sim \text{Meiosis}(\ell_1, \ell_2)$ between $\ell_1, \ell_2$;
        **for** *individuals $i$ in clonal component $C_n$* **do**
            $\boldsymbol{M}[i, \,] \leftarrow \boldsymbol{\ell}_{\text{offspring}}$
**return** $\boldsymbol{M}$

as the truth value for the pairwise relatedness of individuals $j_1, j_2$ in generation $k$.

To diagnose systematic biases in relatedness estimation, we then generate maximum likelihood estimates (MLEs) of the pairwise relatedness parameter $r$ and, where applicable, the switch rate parameter $\kappa$, under both the independence model and HMM using:

- the standard nIBD-to-IBS and (n)IBD-to-allele models predicated on sample allele frequencies;
- the corrected nIBD-to-IBS model based on the sample proportion of nIBD pairs that are IBS at each locus $i$, that is,

$$c_i := \frac{\sum_{1 \le j_1 < j_2 \le n_{\text{individuals}}} \left[ \mathbb{1}\{A_i(j_1,k) = A_i(j_2,k)\} - \mathbb{1}\{F_i(j_1,k) = F_i(j_2,k)\} \right]}{\sum_{1 \le j_1 < j_2 \le n_{\text{individuals}}} \left[ 1 - \mathbb{1}\{F_i(j_1,k) = F_i(j_2,k)\} \right]}.$$

MLEs $(\hat{r}, \hat{\kappa})$ predicated on (n)IBD-to-allele models are generated using the R package `paneljudge` [34], which implements both the HMM and independence model of relatedness. MLEs for the relatedness parameter $r$ predicated on the nIBD-to-IBS model coupled with the independence model of relatedness are computed using a custom R script. We do not generate estimates under the nIBD-to-IBS model coupled with the HMM of relatedness.

Under the ancestrally oblivious HMM of relatedness, each parameter set $(r, \kappa)$ yields a distribution for the realised fraction of IBD markers. Convergence of the realised fraction of IBD markers to the relatedness parameter $r$ occurs in the limit where an infinite number of equidistant markers are sampled along an infinitely-long genome. While we acknowledge the conceptual difference between the pairwise relatedness parameter $r$ and the realised fraction of sites IBD [17, 26], the fact that we model recombination from ancestral principles means that, under our framework, there is no quantity with direct equivalence to $r$.

#### B.5 Summary of simulation parameters

We now examine the effects of simulation parameters on distributions of realised IBD/IBS sharing. A summary of simulation parameters is provided in Table B.1.

##### B.5.1 Effect of $n_{\text{individuals}}$ vs $n_{\text{founders}}$ on background relatedness

We distinguish two sources of relatedness within our simulation framework:

- "Recent" relatedness, attributed to crosses between shared parents from generation one onwards.

| Feature | Parameter | Interpretation | Value |
|---|---|---|---|
| *Markers* | $n_{\text{markers}}$ | Number of equidistant markers distributed across a single chromosome | 24000 |
| | $\rho d$ | Distance between consecutive markers in Morgans | $\approx 0.002$ |
| | $\bar{M} = 2(1 - e^{-\rho d})^{-1}$ | Average number of consecutive markers inherited from the same parent in a cross (reduces to $\bar{M} = 2$ given locus independence) | 1000 |
| *Relationship Graphs* | $n_{\text{individuals}}$ | Number of not necessarily distinct genotypes, each representing an equal number of actual parasites | 100 |
| | $m_{\text{subgraph}}$ | Size of transitive clone/sibling/stranger graphs [27] sampled uniformly at random; interpretable as a MOI upper bound (if subgraphs are viewed as parasite genotypes within a host) or the number of successful, not necessarily distinct genotypes per bite (if subgraphs are viewed as successfuly transmitted sporozoites per bite) | 4 |
| *Founder* | $n_{\text{founders}}$ | Number of unrelated founders | 100 |
| | $y$ | Number of possible alleles at each at marker | 2 |
| | $p_i(q)$ | Probability of each founder $f$ harbouring allele $q \in \{1, \ldots, y\}$ at marker $i$ | $(0.1, 0.9)$ |
| *Inbreeding* | $n_{\text{generations}}$ | Number of discrete, non-overlapping generations of inbreeding [exc. 0] | 10 |
| | $p_{\text{cotransmission}}$ | Probability of sampling parents without replacement from the same constituent subgraph *vs* with replacement from the population as a whole in the previous generation for each sibling/clonal component in a relationship graph | 0.4 |

**Table B.1:** Summary of simulation parameters

- "Background" relatedness, manifest in generation zero parasites, arising from the finite number of founders $n_{\text{founders}}$.

The population size, $n_{\text{individuals}}$ and the number of equifrequent founders $n_{\text{founders}}$ govern the degree of background relatedness at generation zero. At locus $i$, each generation zero parasite is assigned ancestry from a founder $f \in \{1, \ldots, n_{\text{founders}}\}$, with independent assignment across both loci and parasites (Section B.2). The proportion of pairs IBD at locus $i$ in generation zero, $d_i^0$, can thus be written

$$d_i^0 = \frac{X_1^2 + \cdots + X_{n_{\text{founders}}}^2}{n_{\text{individuals}}^2}$$

where

$$(X_1, \ldots, X_{n_{\text{founders}}}) \sim \text{Multinomial}\big(n_{\text{individuals}}, (1/n_{\text{founders}}, \ldots, 1/n_{\text{founders}})\big),$$

and pairs have been sampled with replacement.

In particular, the expected locuswise relatedness at generation zero is given by

$$\mathbb{E}\left[d_i^0\right] = \frac{1}{n_{\text{founders}}}\left[1 - \left(1 - \frac{1}{n_{\text{individuals}}}\right)\left(1 - \frac{1}{n_{\text{founders}}}\right)\right].$$

The degree of background relatedness exhibited by generation zero parasites has a strong bearing on realised IBD distributions in early generations that follow. It also governs the rate at which sibling and half-sibling pedigrees diverge from the expected relatedness 0.5 and 0.25 respectively.

##### B.5.2 Contribution of sibling/clonal subgraphs: $m_{\text{subgraph}}$ vs $n_{\text{individuals}}$

The relative contributions of stranger/sibling/clonal edges in a population-level relationship graph — which are functions of $n_{\text{individuals}}$ and $m_{\text{subgraph}}$ respectively — are important determinants of relatedness. At each generation, we independently sample $n_{\text{individuals}}/m_{\text{subgraph}}$ subgraphs of size $m_{\text{subgraph}}$ (uniformly at random, over the set of all allowable transitive graphs [27, 35]) and knit them together with stranger edges. As such, all between-subgraph comparisons — which constitute proportion $(n_{\text{individuals}} - m_{\text{subgraph}})/n_{\text{individuals}}$ of all parasite pairs (with replacement) — necessarily correspond to stranger edges. Of the remaining proportion of pairwise comparisons, $m_{\text{subgraph}}/n_{\text{individuals}}$, the weighting of sibling/clonal edges is contingent on the underlying uniform distribution over all allowable transitive graphs of size $m_{\text{subgraph}}$; the case $m_{\text{subgraph}} = 4$ is shown in Figure B.2A.

Population-level relationship graphs enriched for sibling and clonal edges naturally yield rapid growth in relatedness over successive generations of inbreeding. For parasite pairs connected by a stranger edge in generation $k$, parents are sampled independently from generation $(k -$

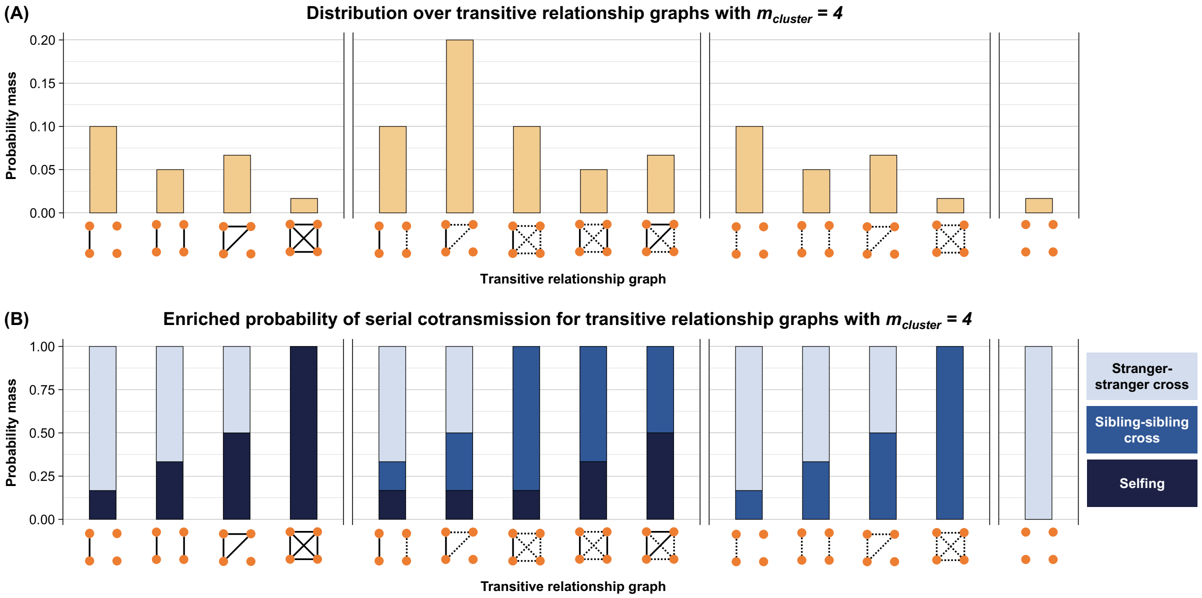

**Figure B.2:** (A) Probability masses for transitive relationship graphs with sibling/clone/stranger edges [27] in the case $m_{\text{subgraph}} = 4$, under a uniform distribution over all allowable graphs. Clonal and sibling edges are indicated with solid and dashed lines, respectively. (B) Probability of stanger-stranger crosses vs sibling-sibling crosses vs selfing when two parents are sampled uniformly at random *without* replacement from each allowable relationship graph with $m_{\text{subgraph}} = 4$.

1). Half-sibling, sibling or clonal relationships between individuals connected by a stranger edge in generation $k$ can therefore arise due to random sampling from the finite population in generation $(k-1)$, and are a direct function of clonal structure in generation $(k-1)$. However, we would expect half-sibling, sibling and clonal relationships in generation $k$ to be driven primarily by sibling/clonal edges within the population-level relatedness graph for generation $k$ itself — particularly in early generations with limited clonal substructure.

Under our framework, we recover a mixture distribution for pairwise IBD sharing $R(j_1, j_2, k)$ in generation $k$ — with a dominant component attributable to parasites separated by at least two generations and smaller components attributable to clones, siblings and half-siblings. The size $m_{\text{subgraph}}$ of uniformly-sampled transitive graphs with sibling/clonal/stranger edges [27, 35], relative to the parasite population size $n_{\text{individuals}}$, governs the relative weightings of the half-sibling, sibling and clonal components to the mixture distribution — and consequently, the rate at which relatedness is accrued over successive generations.

##### B.5.3 Breeding between closely-related parents: $p_{\mathbf{cotransmission}}$ vs $m_{\mathbf{subgraph}}$

For each sibling/clonal component within the population-graph of generation $k$, with probability $p_{\text{cotransmission}}$, we have sampled parents without replacement from a subgraph of size $m_{\text{subgraph}}$ in population graph of generation $(k-1)$. Under our simulation structure, each subgraph is sampled uniformly at random over the set of all allowable transitive relationship graphs of size $m_{\text{subgraph}}$. The probability of a stranger-stranger cross vs a sibling-sibling cross vs selfing under this regime is therefore a function of $m_{\text{subgraph}}$, with the case $m_{\text{subgraph}} = 4$ illustrated in Figure B.2B. The underlying rationale of this sampling scheme is to yield an enriched probability of sibling-sibling crosses, as a proxy for serial cotransmission [24]; sampling from the constitient subgraphs *without* replacement augments the contribution of sibling-sibling crosses relative to selfing.

Setting $p_{\text{cotransmission}} = 0$ — whereby the sampling of parents is predicated on stochastic drift only (Section B.5.2) — yields a 'blindspot' in our simulated distributions of realised relatedness: in contrast to empirical estimates, we recover negligibly few *non-clonal* parasite pairs with IBD sharing exceeding 60%, signifying crosses between related parents. To recapitulate the smattering of parasite pairs with IBD sharing in the range 60% to 90% seen in empirical estimates, we must inflate the probability of breeding between closely-related parental lineages, consistent with evidence of extensive cotransmission [28]. We modulate $p_{\text{contransmission}} > 0$ and exploit substructure within simulated population-level relatedness graphs to mitigate this blindspot. While we have not explicitly embedded a mechanistic transmission model within our framework, subgraphs derived from parental lineages sampled from the population at large can be thought to encompass superinfection; while subgraphs derived from parental lineages sampled from a previous subgraph can be thought to correspond to cotransmission.

We note, however, that the augmented probability of sibling-sibling crosses under $p_{\text{cotransmission}} > 0$ yields elevated IBD sharing (in the vicinity of 60% to 90%) for a relatively small proportion of parasite pairs: it primarily affects within-component comparisons for each sibling/clonal component in the accompanying population-level relationship graph, with an upper bound $m_{\text{subgraph}}$ on the size of each sibling/clonal component. Between-cluster comparisons are largely modulated by stochastic drift, as discussed in Section B.5.2. Reverting to the mixture distribution interpretation of simulated pairwise IBD delineated in Section B.5.2, setting $p_{\text{contransmission}} > 0$ principally has the effect of broadening the sibling component, to include a small proportion of pairs reflecting breeding between recently-related parents.

##### B.5.4 Linkage structure and spanning (n)IBD segments: $n_{\mathbf{markers}}$ vs $\bar{M}$

The mean length of consecutive (n)IBD markers $\bar{M}$ in a single meiosis governs the degree of dependence between loci, and consequently, the amount of linkage structure exhibited by related parasite pairs. For this linkage structure to emerge, however, the $n_{\text{markers}}$ of interest must span sufficiently many segments of (n)IBD markers; and each segment of (n)IBD markers must be sufficiently long. As a function of the pairwise relatedness parameter $r^{(k,\ell)}$, the interplay between $\bar{M}$ and $n_{\text{markers}}$ also governs the variance of the realised fraction of (polymorphic) sites IBD — which we treat as our truth value to diagnose biases in relatedness estimation.

We adopt a physical argument to recover $\bar{M}$ as a function of the marker count $n_{\text{markers}}$: for some number if equidistant markers $n_{\text{markers}}$, distributed across a chromosome of length $L$ Morgans,

$$\bar{M}(n_{\text{markers}}) = \frac{2}{1 - e^{-L/n_{\text{markers}}}}.$$

##### B.5.5 Minor allele frequency spectra: $\boldsymbol{p}_i$

At each locus $i = 1, \ldots, n_{\text{markers}}$, the count $n_{\text{founders}} \cdot \theta_{\text{founder}}(q, i)$ of each allele $q \in \{1, \ldots, y\}$ in the founder population (size $n_{\text{founders}}$) follows a multinomial distribution

$$n_{\text{founders}} \cdot \big(\theta_{\text{founder}}(1, i), \ldots, \theta_{\text{founder}}(y, i)\big) \sim \text{Multinomial}\big(n_{\text{founders}}, (p_i(1), \ldots, p_i(y))\big)$$

where $p_i(q)$ is the probability of assigning allele $q$ to a founder at locus $i$.

In the present manuscript, we consider only biallelic loci, i.e., we fix $y = 2$. The parameter $p_i$ governing founder sample allele frequencies $\theta_{\text{founder}}$ has a strong bearing on minor allele frequency (MAF) spectra observed across successive generations. In the infinite generation limit, we would expect an allele $q_i \in \{1, 2\}$ to reach fixation at each locus $i$, akin to the canonical Wright-Fisher model of stochastic drift [15]. As in the Wright-Fisher model, however, it is not necessarily the case that the *major* allele in the founder population reaches fixation. As such, in a transient setting, we can either observe allele frequencies becoming increasingly balanced over successive generations (as the minor allele grows in frequency); or increasingly unbalanced (as the major allele grows in frequency). For the most part, however we expect a reduction in genetic diversity as crosses are repeatedly assigned major alleles.

To recover positively-skewed biallelic MAF spectra with zero mode in the order of 10 generations, we require $\min\{p_i(1), p_i(2)\} \approx 0.1$, translating to unbalanced founder allele frequencies. Under this setting, however, we find that allele frequencies become increasingly balanced over successive generations for a subset of markers.

### Appendix C

### Glossary of terms

**$\overline{\text{IBD}}$** proportion of pairs of sampled parasites (with replacement, i.e., including self-self comparisons) that are IBS at a given locus.

**$\overline{\text{IBS}}$** proportion of pairs of sampled parasites (with replacement, i.e., including self-self comparisons) that are IBS at a given locus.

**(n)IBD-to-observation model** inclusive of nIBD-to-allele, IBD-to-allele and nIBD-to-IBS models, under the assumed absence of genotyping error (whereby IBD necessarily implies IBS, i.e., IBD-to-IBS=1).

**background relatedness** under the simulation model, relatedness structure in generation zero parasites.

**genotype** a specific realisation of the genome, which is a random variable distributed according to some ancestral process; in this study, we consider the genotype to be a sequence over all polymorphisms in the genome (elsewhere, its definition extends to subsets).

**inbreeding** recombination between genetically different but related parasites.

**individual** a parasite genotype.

**meiotic siblings** siblings derived from the same oocyst, i.e., siblings that are complements of one another and thus not independent.

**outbreeding** recombination between genetically unrelated parasites.

**realised relatedness** the fraction of polymorphic markers within the sample that are IBD for a given parasite pair.

**recent relatedness** under the simulation model, relatedness structure resulting from inbreeding from generation one onwards.

**relatedness structure** the locuswise partition of individuals in a parasite population or sample into transitive IBD clusters.

**sample** a collection of individuals, i.e., a collection of parasite genotypes.

**selfing** recombination between genetically identical parasites.

**strangers** a pair of individuals separated by two or more generations of recombination.
