## Supplementary material for "Systematic bias in malaria parasite relatedness estimation": Table S1

| ID | Accession | Country | Year |
|---|---|---|---|
| G1P001 | | Guyana | 2020 |
| G1P002 | | Guyana | 2020 |
| G1P005 | | Guyana | 2020 |
| G1P007 | | Guyana | 2020 |
| G1P009 | | Guyana | 2020 |
| G1P010 | | Guyana | 2020 |
| G1P013 | | Guyana | 2020 |
| G1P015 | | Guyana | 2020 |
| G1P024 | | Guyana | 2020 |
| G1P025 | | Guyana | 2020 |
| G1P026 | | Guyana | 2020 |
| G1P028 | | Guyana | 2020 |
| G1P029 | | Guyana | 2020 |
| G1P030 | | Guyana | |
| G1P031 | | Guyana | 2020 |
| G1P032 | | Guyana | 2020 |
| G1P033 | | Guyana | 2020 |
| G1P034 | | Guyana | |
| G1P035 | | Guyana | 2020 |
| G1P036 | | Guyana | |
| G1P051 | | Guyana | 2020 |
| G1P073 | | Guyana | 2020 |
| G1P076 | | Guyana | 2020 |
| G1P080 | | Guyana | 2020 |
| G1P091 | | Guyana | 2020 |
| G1P100 | | Guyana | 2020 |
| G1P101 | | Guyana | 2020 |
| G1P105 | | Guyana | 2020 |
| G1P111 | | Guyana | 2020 |
| G1P112 | | Guyana | 2020 |
| G1P115 | | Guyana | 2020 |
| G1P120 | | Guyana | 2020 |
| G1P156 | | Guyana | 2020 |
| G1P157 | | Guyana | 2020 |
| G1P158 | | Guyana | 2020 |
| G1P160 | | Guyana | 2020 |
| G1P161 | | Guyana | 2020 |
| G1P162 | | Guyana | 2020 |
| G1P166 | | Guyana | 2020 |
| G1P167 | | Guyana | 2020 |
| G1P168 | | Guyana | 2020 |
| G1P170 | | Guyana | 2020 |
| G1P171 | | Guyana | 2020 |
| G1P172 | | Guyana | 2020 |
| G1P173 | | Guyana | 2020 |
| G1P175 | | Guyana | 2020 |
| G4G007 | | Guyana | 2020 |
| G4G008 | | Guyana | 2020 |
| G4G010 | | Guyana | 2020 |
| G4G013 | | Guyana | 2020 |
| G4G015 | | Guyana | 2020 |
| G4G016 | | Guyana | 2020 |
| G4G024 | | Guyana | 2020 |

| | | | |
|---|---|---|---|
| G4G025 | | Guyana | 2020 |
| G4G026 | | Guyana | 2020 |
| G4G027 | | Guyana | 2020 |
| G4G028 | | Guyana | 2020 |
| G4G035 | | Guyana | 2020 |
| G4G036 | | Guyana | 2020 |
| G4G041 | | Guyana | 2020 |
| G4G043 | | Guyana | 2020 |
| G4G044 | | Guyana | 2020 |
| G4G045 | | Guyana | 2020 |
| G4G047 | | Guyana | 2020 |
| G4G048 | | Guyana | 2020 |
| G4G049 | | Guyana | 2020 |
| G4G050 | | Guyana | 2020 |
| G4G055 | | Guyana | 2020 |
| G4G057 | | Guyana | 2020 |
| G4G060 | | Guyana | 2020 |
| G4G061 | | Guyana | 2020 |
| G4G063 | | Guyana | 2020 |
| G4G064 | | Guyana | 2020 |
| G4G067 | | Guyana | 2020 |
| G4G069 | | Guyana | 2020 |
| G4G070 | | Guyana | |
| G4G073 | | Guyana | 2020 |
| G4G074 | | Guyana | 2020 |
| G4G089 | | Guyana | 2020 |
| G4G093 | | Guyana | 2020 |
| G4G094 | | Guyana | 2020 |
| G4G096 | | Guyana | 2020 |
| G4G098 | | Guyana | 2020 |
| G4G100 | | Guyana | 2020 |
| G4G103 | | Guyana | 2020 |
| G4G105 | | Guyana | 2020 |
| G4G114 | | Guyana | 2020 |
| G4G118 | | Guyana | 2020 |
| G4G128 | | Guyana | 2020 |
| G4G130 | | Guyana | 2020 |
| G4G131 | | Guyana | 2020 |
| G4G134 | | Guyana | 2020 |
| G4G138 | | Guyana | 2020 |
| G4G141 | | Guyana | 2020 |
| G4G154 | | Guyana | 2020 |
| G4G156 | | Guyana | 2020 |
| G4G157 | | Guyana | 2020 |
| G4G161 | | Guyana | 2020 |
| G4G162 | | Guyana | 2020 |
| G4G164 | | Guyana | 2020 |
| G4G167 | | Guyana | 2020 |
| G4G172 | | Guyana | 2020 |
| G4G174 | | Guyana | 2020 |
| G4G177 | | Guyana | 2020 |
| G4G179 | | Guyana | 2020 |
| G4G180 | | Guyana | 2020 |
| G4G182 | | Guyana | 2020 |

| | | | |
|---|---|---|---|
| G4G185 | | Guyana | 2020 |
| G4G186 | | Guyana | 2020 |
| G4G190 | | Guyana | 2020 |
| G4G191 | | Guyana | 2020 |
| G4G192 | | Guyana | 2020 |
| G4G193 | | Guyana | 2020 |
| G4G195 | | Guyana | 2020 |
| G4G198 | | Guyana | 2020 |
| G4G199 | | Guyana | 2020 |
| G4G201 | | Guyana | 2020 |
| G4G202 | | Guyana | 2020 |
| G4G205 | | Guyana | 2020 |
| G4G209 | | Guyana | 2020 |
| G4G213 | | Guyana | 2020 |
| G4G215 | | Guyana | 2020 |
| G4G223 | | Guyana | 2020 |
| G4G225 | | Guyana | 2020 |
| G4G237 | | Guyana | 2020 |
| G4G239 | | Guyana | 2020 |
| G4G241 | | Guyana | 2020 |
| G4G242 | | Guyana | 2020 |
| G4G246 | | Guyana | 2020 |
| G4G253 | | Guyana | 2020 |
| G4G255 | | Guyana | 2020 |
| G4G256 | | Guyana | 2020 |
| G4G257 | | Guyana | 2020 |
| G4G258 | | Guyana | 2020 |
| G4G263 | | Guyana | 2020 |
| G4G267 | | Guyana | 2020 |
| G4G270 | | Guyana | 2020 |
| G4G271 | | Guyana | 2020 |
| G4G274 | | Guyana | 2020 |
| G4G278 | | Guyana | 2020 |
| G4G281 | | Guyana | 2020 |
| G4G282 | | Guyana | 2020 |
| G4G284 | | Guyana | 2020 |
| G4G288 | | Guyana | 2020 |
| G4G305 | | Guyana | 2020 |
| G4G306 | | Guyana | 2020 |
| G4G318 | | Guyana | 2020 |
| G4G323 | | Guyana | 2020 |
| G4G324 | | Guyana | 2020 |
| G4G327 | | Guyana | 2020 |
| G4G330 | | Guyana | 2020 |
| G4G331 | | Guyana | 2020 |
| G4G339 | | Guyana | 2020 |
| G4G342 | | Guyana | 2020 |
| G4G346 | | Guyana | 2020 |
| G4G351 | | Guyana | 2020 |
| G4G353 | | Guyana | 2020 |
| G4G356 | | Guyana | 2020 |
| G4G357 | | Guyana | 2020 |
| G4G359 | | Guyana | 2020 |
| G4G360 | | Guyana | 2020 |

| | | | |
|---|---|---|---|
| G4G362 | | Guyana | 2020 |
| G4G371 | | Guyana | 2020 |
| G4G372 | | Guyana | 2020 |
| G4G373 | | Guyana | 2020 |
| G4G380 | | Guyana | 2020 |
| G4G384 | | Guyana | 2020 |
| G4G392 | | Guyana | 2020 |
| G4G396 | | Guyana | 2020 |
| G4G404 | | Guyana | 2020 |
| G4G408 | | Guyana | 2020 |
| G4G409 | | Guyana | 2020 |
| G4G410 | | Guyana | 2020 |
| G4G411 | | Guyana | 2020 |
| G4G414 | | Guyana | 2020 |
| G4G422 | | Guyana | 2020 |
| G4G427 | | Guyana | 2020 |
| G4G429 | | Guyana | 2020 |
| G4G434 | | Guyana | 2020 |
| G4G439 | | Guyana | 2020 |
| G4G443 | | Guyana | 2020 |
| G4G444 | | Guyana | 2020 |
| G4G446 | | Guyana | 2020 |
| G4G447 | | Guyana | 2020 |
| G4G450 | | Guyana | 2020 |
| G4G452 | | Guyana | 2020 |
| G4G463 | | Guyana | 2020 |
| G4G469 | | Guyana | 2020 |
| G7B002 | | Guyana | 2020 |
| G7B003 | | Guyana | 2020 |
| G7B005 | | Guyana | 2020 |
| G7B008 | | Guyana | 2020 |
| G7B017 | | Guyana | 2020 |
| G7B018 | | Guyana | 2020 |
| G7B021 | | Guyana | 2020 |
| G7B028 | | Guyana | 2020 |
| G7B035 | | Guyana | 2020 |
| G7B037 | | Guyana | 2020 |
| G7B041 | | Guyana | 2020 |
| G7B043 | | Guyana | 2020 |
| G7B051 | | Guyana | 2020 |
| G7B054 | | Guyana | 2020 |
| G7B056 | | Guyana | 2020 |
| G7B057 | | Guyana | 2020 |
| G7B066 | | Guyana | 2020 |
| G7B069 | | Guyana | 2020 |
| G7B071 | | Guyana | 2020 |
| G7B072 | | Guyana | 2020 |
| G7B073 | | Guyana | 2020 |
| G7B074 | | Guyana | 2020 |
| G7B081 | | Guyana | 2020 |
| G7B082 | | Guyana | 2020 |
| G7B084 | | Guyana | 2020 |
| G7B088 | | Guyana | 2020 |
| G7B095 | | Guyana | 2020 |

| | | | |
|---|---|---|---|
| G7B098 | | Guyana | 2020 |
| G7B100 | | Guyana | 2020 |
| G7B101 | | Guyana | 2020 |
| G7B107 | | Guyana | 2020 |
| G7B108 | | Guyana | 2020 |
| G7B109 | | Guyana | 2020 |
| G7B117 | | Guyana | 2020 |
| G7B127 | | Guyana | 2020 |
| G7B130 | | Guyana | 2020 |
| G7B139 | | Guyana | 2020 |
| G7B140 | | Guyana | 2020 |
| G7B142 | | Guyana | 2020 |
| G7B145 | | Guyana | 2020 |
| G7B146 | | Guyana | 2020 |
| G7B147 | | Guyana | 2020 |
| G7B148 | | Guyana | 2020 |
| G7B149 | | Guyana | 2020 |
| G7B151 | | Guyana | 2020 |
| G7B155 | | Guyana | 2020 |
| G7B164 | | Guyana | 2020 |
| G7B165 | | Guyana | 2020 |
| G7B167 | | Guyana | 2020 |
| G7B174 | | Guyana | 2020 |
| G7B178 | | Guyana | 2020 |
| G7B187 | | Guyana | 2020 |
| G7B189 | | Guyana | 2020 |
| G7B193 | | Guyana | 2020 |
| G7B194 | | Guyana | 2020 |
| G7B195 | | Guyana | 2020 |
| G7B197 | | Guyana | 2020 |
| G7B199 | | Guyana | 2020 |
| G7B210 | | Guyana | 2020 |
| G7B217 | | Guyana | 2020 |
| G7B225 | | Guyana | 2020 |
| GUY0123 | | Guyana | 2017 |
| GUY0154 | | Guyana | 2017 |
| GUY0159 | | Guyana | 2017 |
| GUY0166 | | Guyana | 2017 |
| GUY0174 | | Guyana | 2017 |
| T104 | | Guyana | 2016 |
| T118 | | Guyana | 2016 |
| T133 | | Guyana | 2016 |
| T145_swga | | Guyana | 2016 |
| T158 | | Guyana | 2016 |
| T159 | | Guyana | 2016 |
| T211 | | Guyana | 2016 |
| T215 | | Guyana | 2016 |
| T227 | | Guyana | 2016 |
| T230 | | Guyana | 2016 |
| T240 | | Guyana | 2016 |
| T285 | | Guyana | 2016 |
| T301 | | Guyana | 2016 |
| T359 | | Guyana | 2016 |
| T373 | | Guyana | 2016 |

| | | | |
|---|---|---|---|
| T377 | | Guyana | 2016 |
| T387 | | Guyana | 2016 |
| T392 | | Guyana | 2016 |
| T482 | | Guyana | 2016 |
| T497 | | Guyana | 2016 |
| T640 | | Guyana | 2016 |
| T679 | | Guyana | 2016 |
| T824 | | Guyana | 2016 |
| T827 | | Guyana | 2016 |
| PW0069-C | ERR1818180 | Colombia | 2014-17 |
| PW0070-C | ERR1818181 | Colombia | 2014-17 |
| PW0008-C | ERR039903 | Colombia | 1993-2007 |
| PW0002-C | ERR039930 | Colombia | 1993-2007 |
| PW0009-C | ERR039986 | Colombia | 1993-2007 |
| PW0016-C | ERR039988 | Colombia | 1993-2007 |
| PW0007-C | ERR042222 | Colombia | 1993-2007 |
| PW0017-C | ERR042223 | Colombia | 1993-2007 |
| PW0012-C | ERR042224 | Colombia | 1993-2007 |
| PW0004-C | ERR042226 | Colombia | 1993-2007 |
| PW0003-C | ERR042227 | Colombia | 1993-2007 |
| PW0001-C | ERR042228 | Colombia | 1993-2007 |
| PW0005-C | ERR042229 | Colombia | 1993-2007 |
| PW0006-C | ERR042230 | Colombia | 1993-2007 |
| PW0013-C | ERR042231 | Colombia | 1993-2007 |
| PW0015-C | ERR042232 | Colombia | 1993-2007 |
| PW0014-C | ERR042233 | Colombia | 1993-2007 |
| PW0053-C | ERR1818164 | Colombia | 2014-17 |
| PW0057-C | ERR1818168 | Colombia | 2014-17 |
| PW0067-C | ERR1818178 | Colombia | 2014-17 |
| PW0105-C | ERR1911267 | Colombia | 2014-17 |
| SPT26227 | ERR2496598 | Colombia | 2014-17 |
| PW0073-C | ERR1818184 | Colombia | 2014-17 |
| PW0088-C | ERR1911250 | Colombia | 2014-17 |
| SPT26335 | ERR2496593 | Colombia | 2014-17 |
| SPT26336 | ERR2496568 | Colombia | 2014-17 |
| SPT26229 | ERR2496572 | Colombia | 2014-17 |
| SPT26248 | ERR2496548 | Colombia | 2014-17 |
